# High resolution reconstruction of a Jumbo bacteriophage infecting capsulated bacteria using hyperbranched tail fibers

**DOI:** 10.1101/2022.03.15.484430

**Authors:** Ruochen Ouyang, Ana Rita Costa, C. Keith Cassidy, Aleksandra Otwinowska, Vera C. J. Williams, Agnieszka Latka, Phill J. Stansfeld, Zusanna Drulis-Kawa, Yves Briers, Daan M. Pelt, Stan J. J. Brouns, Ariane Briegel

## Abstract

The *Klebsiella* jumbo myophage ϕKp24 displays an unusually complex arrangement of tail fibers interacting with a host cell. In this study, we combined cryo-electron microscopy methods, protein structure prediction methods, molecular simulations, and machine learning approaches to explore the capsid, tail, and tail fibers of this phage at high resolution. We determined the structure of the capsid and tail at 4.3Å and 4.1Å resolution. We observed that the tail fibers were highly branched and rearranged dramatically upon cell surface attachment. This complex configuration involves fourteen putative tail fibers with depolymerase activity that provide ϕKp24 with the ability to infect a broad panel of capsular polysaccharide (CPS) types of *Klebsiella pneumoniae*. Taken together, our study provides structural and functional insight into how ϕKp24 adapts to the highly variable surfaces of capsulated bacterial pathogens, which will be useful for the development of phage therapy approaches against pan-drug resistant *K. pneumoniae* strains.

## INTRODUCTION

Disease-causing bacteria pose an ever-increasing threat to human health. While many bacterial infections can be effectively cured by antibiotics, antimicrobial resistance (AMR) is increasingly resulting in ineffective treatments. A recent study reported 3.57 million deaths associated with AMR in six leading pathogens in 2019 (Murray et al., 2022). Among these is *Klebsiella pneumoniae*, a pathogen recognized by the World Health Organization as a priority for the development of new antibiotics (Tacconelli et al., 2018). *K. pneumoniae* can cause pneumonia, urinary tract infection, bacteremia, and other infectious diseases in humans, especially individuals with weakened immunity (Ryan and Ray, 2004). The vast majority of *K. pneumoniae* clinical isolates express a pronounced capsule that is generally considered as an important virulence factor mediating protection from the host’s immune system (Ko, 2017). There are over a hundred genetically distinct capsular locus types, of which 77 well-characterized chemical structures are used in serotyping (K types) (Follador et al., 2016).

Even though multidrug-resistant *K. pneumoniae* are resistant to standard-of-care antibiotics, they remain susceptible to bacteriophage infection. Bacteriophages, or phages for short, are viruses that infect bacteria. Klebsiella-specific phages can successfully infect and kill their natural host; however, they are typically highly strain-specific due to the variable capsular polysaccharides (CPS) of this species, which act as a primary phage receptor. Capsule-dependent Klebsiella phages, including jumbo phages (Hendrix, 2009; Yuan and Gao, 2017) ΦK64-1 or vB_KleM-RaK2 (Pan et al., 2017; Šimoliūnas et al., 2013), are equipped with receptor binding proteins (RBPs) containing CPS degrading enzyme domains (coined depolymerases) that enable successful phage adsorption and infection (Latka et al., 2019) of a limited number of capsular types. RBPs can adopt a tail fiber or tailspike shape. For simplicity, we henceforth use the term ‘tail fiber’.

The recently described vB_KpM_FBKp24 (ϕKp24) jumbo myophage encodes at least nine tail fibers containing different depolymerase domains, which suggests an expanded host range. Genomic analysis revealed that ϕKp24 has very limited similarity to any other known phage (Bonilla et al., 2021) and therefore constitutes a new family termed *vanleeuwenhoekviridae* (ICTV classification). Furthermore, transmission electron microscopy revealed a unique complex structure of tail fibers at the baseplate. However, more detailed insight into the structure and function of this unique set of tail fibers is currently lacking.

To gain insight into the unusual structure of ϕKp24, we analyzed its capsid, tail, and tail fibers using different structural methods. We generated atomic models for the highly ordered phage capsid and tail using cryo-electron microscopy (cryo-EM) single particle analysis (SPA) combined with AlphaFold2 (Cramer, 2021) protein structure predictions and molecular dynamics (MD) simulations. The data revealed unusual features of the capsid, most notably the composition of the entire capsid by a single MCP, and the presence of a tube in the center of the hexagons. We obtained insight into the structure of the flexible and disordered tail fibers using cryo-electron tomography (cryo-ET). This analysis was aided by machine learning approaches to train a neural network to automatically track the complex tail fibers for quantitative analysis of the tomography data. Our analysis revealed a pronounced rearrangement of the tail fibers during the infection process. In addition, we tested the infectivity of ϕKp24 using a K serotype collection of *K. pneumoniae* strains, showing the unusually broad panel of CPS types targeted by the tail fibers.

We combined the structural investigation of ϕKp24 with an analysis of the tail fibers based on sequence prediction and binding assays against a Klebsiella serotype library in combination with a structure-function analysis of the tail fibers to get insights in their complex organization. The combination of structural, computational, and laboratory techniques allowed us to gain in-depth insight into the structure and function of this distinct bacteriophage.

Furthermore, the insights gained from this novel combination of diverse methods provide an essential basis for potential clinical applications. Recently, phage therapy has been successfully applied to treat an infection caused by a pan-drug resistant *K. pneumoniae* strain (Eskenazi et al., 2022). The extended host range of ϕKp24 determined here may make it an attractive candidate for the development of phage therapy.

## RESULTS

### Capsid structure of phage ϕKp24

To study the structure of the ϕKp24 capsid and tail, we imaged the phage using cryo-EM (Figure. S1A, data collection parameters in Figure. S1B). In our dataset, we find three variants of the capsid: full capsids filled with DNA, empty capsids, and partially DNA-filled capsids. We reconstructed the full (EMD-14356, Figure. S2A) and empty (EMD-13862, Figure. 1A) capsid structures of ϕKp24 with Relion (Scheres, 2012) to 4.7 Å and 4.3 Å resolution after polishing, respectively, using the “gold-standard” FSC_0.143_ criterion (Rosenthal and Henderson, 2003) (Figures. S2B and S3B). Since the empty and full capsid reconstructions and superimpose well (Figure. S4), we focused our analysis on the higher-resolution reconstruction of the empty capsid. The empty capsid (Figure. 1A) has a diameter of 145 nm from vertex to vertex, 120 nm along the 2-fold symmetry axis, and follows a T = 27 (h = 3; k = 3; T = ℎ^2^ + *k*^2^ + ℎ*k* ) triangulation symmetry with a planar outline. (Figure. 1B).

**Figure 1.**
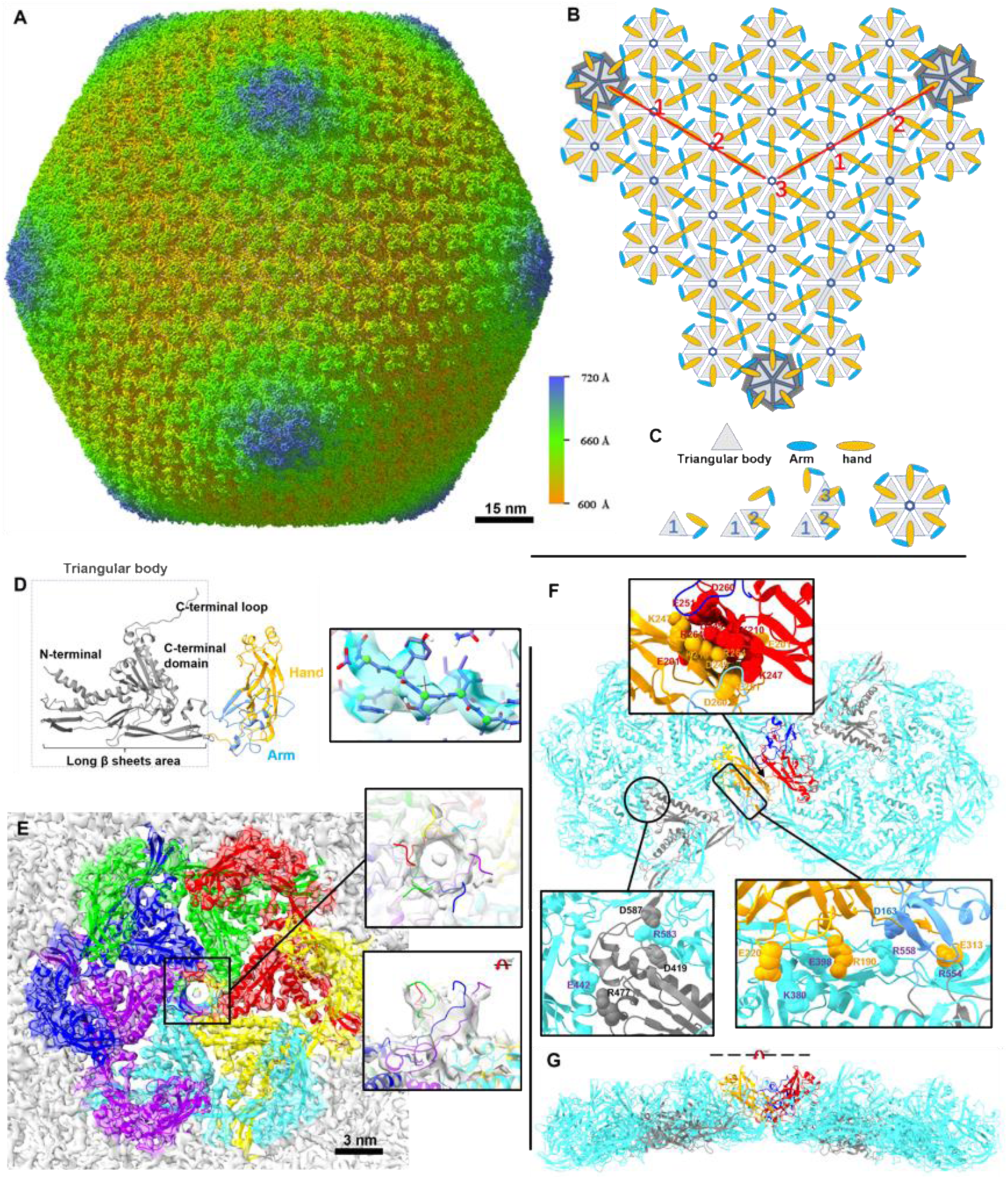
Reconstruction and organization of the empty capsid of phage ϕKp24, MCP, hexamer arrangement, and major capsid protein. (A) The dsDNA-empty capsid (EMD-13862) of phage ϕKp24 was reconstructed at 4.3 Å resolution using 19,193 particles. The displayed structure is colored radially based on the distance from the capsid center using the color bar on the lower right. (B) Schematic illustrating the ϕKp24 capsid organization. The capsid follows a T = 27 symmetry. The h and k volumes in triangulation symmetry calculation are highlighted in red (h is the number of units in straight line toward next pentamer, k is the number of units shifted in either side to reach the next pentamer (https://viralzone.expasy.org/8577)). (C)Organization of one hexamer. A hexamer can interact with other hexamers or a pentamer via arms (blue) or hands (orange) which likely have a stabilizing function similar to decoration proteins present in other phage structures. (D) Ribbon diagram of gp372 from phage ϕKp24 generated by AlphaFold2. The model consists of three parts colored corresponding to the monomer of a hexamer in Figure. 1C. The right panel shows the Cryo-EM density map and an atomic model of gp372. (E) Top view of an hexameric capsomere. The six monomers are colored individually. At the center of the hexamer, a pore is formed by six long C-terminal loops twisting together (enlarged rectangular box, right upper panel). The right lower panel shows the pore after rotation of the upper image by 90°. (F) Two hexamers with highlighted inter-capsomere interactions. Here, two gp372 hand regions interact with each other via a large cluster of charged residues, including E201, K210, K247, D249, E251, D260, and R264 (enlarged rectangular box, upper panel). The MCP triangular bodies within the same hexamer interact using complementary salt bridges between residue R477 and E442, residue R583 and residues D419 and D587 of the clockwise neighbor (enlarged rectangular box, lower left panel). The triangular body also interacts with the hand region through a series of salt bridges, including K380/E220, E398/R190, R554/E313, and R558/D163 (enlarged rectangular box, lower right panel). Hydrogen bonds were analyzed using the UCSF ChimeraX with relaxed distance and angle criteria (0.4 Å and 20° tolerance, respectively). (G) Side view of hexamer-hexamer interactions after rotation of the upper image by 90°.

Similar to other phage capsids (Duda and Teschke, 2019; Suhanovsky and Teschke, 2015), the capsid of ϕKp24 is built up by a lattice of hexamers and pentamers in each facet (Figure. 1B). Unlike other capsids, the hexamers and pentamers are made up of copies of the same major capsid protein (MCP, gp372, accession no QQV92002). The capsid contains a total of 260 hexamers (13 per facet) and 11 pentamers (one vertex is occupied by the portal complex), which represent 1,615 copies of the major capsid protein (MCP) (Yamada et al., 2010). The MCPs are assembled into an icosahedral shell that encloses the phage genome. In the case of ϕKp24, the predicted MCP is gp372 (Figure. 1D, left).

Overall, we reconstructed the capsid of phage Kp24 at the highest resolution of a Jumbo phage to date, revealing a T = 27 symmetry build up from multiple copies of a single protein.

### Structure of the major capsid protein of ϕKp24

Gp372 contains 597 amino acids and is to our knowledge the largest MCP currently described in any phage. Its structure was predicted and modeled using AlphaFold2 (Cramer, 2021) (Figure. 1D, left). It consists of three major parts: a “triangular body” (Figure. 1C, gray triangle, and Figure. 1D, dotted box, residues 28-88, 314-597); an “arm” (Figure. 1C, blue oval, and Figure. 1D, blue, residues 89-166); and a “hand” (Figure. 1C, orange oval, and Figure. 1D, orange, residues 167-313).

The triangular body shows a core fold similar to the MCP gp5 of Enterobacteria phage HK97 (Gan et al., 2006; Helgstrand et al., 2003). However, there are some notable differences between the MCP of ϕKp24 and HK97. Gp372 of ϕKp24 contains a long C-terminal loop (a component of the pore) that is absent in HK97. Additionally, the N-terminal and C-terminal domains are folded differently: (1) the A domain of HK97 gp5 monomer has four central, mostly anti-parallel β-sheets and two helices (Helgstrand *et al*., 2003). In contrast, there are five anti-parallel β-sheets and four helices of different length in the corresponding domain of gp372 (C-terminal domain); (2) the N-terminal arm of HK97 gp5 monomer is in a mostly unstructured conformation, while the N-terminal of gp372 folds as a long helix (Figure. S5).

The arm consists of two short anti-parallel β-sheets, three short helices, and one short β-sheet close to the “hand”. The most prominent feature of the hand is two layers of β-sheets, with each layer containing three anti-parallel β-sheets. Additionally, there are two short helices at the tip of “hand”.

To further refine the gp372 structure according to our 3D cryo-EM data and to determine putative interactions between MCPs in the native capsid assembly, we next reconstructed models of the capsid hexamer and pentamer assemblies, including the nearest neighboring hexamers (Figure. S6). We then flexibly refined these models to our 4.3Å EM map using ISOLDE and molecular dynamics flexible fitting (MDFF) (McGreevy et al., 2016). Although the map resolution is not sufficient for accurate placement of all residue sidechains, we can still observe prominent densities throughout the gp372 structure corresponding to likely side chain conformations (Figure. 1D, right), allowing for an assessment of key residue-residue interactions between neighboring MCP monomers that are likely crucial for capsid stability.

### Inter-molecular interactions within and between hexamer units

After analyzing the inter-MCP contacts of the hexamer, we assembled a list of putative strong interactions (Figure. S7). The triangular body of the MCPs interacts primarily with the triangular bodies of the MCPs directly adjacent and within the same hexamer (Figure. 1F). These interactions include a series of complimentary salt bridges between residue R477 and E442 on the clockwise neighbor, as well as residue R583 and residues D419 and D587 on the clockwise neighbor. Additionally, the triangular body also interacts strongly with the hand region of its clockwise neighbor through another series of salt bridges (Figure. 1F), including K380/E220, E398/R190, R554/E313, and R558/D163. In addition to these interactions formed between MCPs within the same hexamer, the hand region also mediates strong interactions with the hand region of an MCP from an adjacent hexamer (Figure. 1F). These involve a large cluster of charged residues, including E201, K210, K247, D249, E251, D260, and R264, which form complimentary interactions. Finally, compared to other known phage capsid structures (Kamiya et al., 2021; Wikoff et al., 2000), the MCP of ϕKp24 displayed a structural feature that has, to our knowledge, not been previously reported: the long C-terminal loop (residues Q582-S597) participates in the formation of a pore with an inner diameter of approximately 2 nm (Figure. 1E, right upper and lower panels). Six of these long loops twist together to form the pore in the center of a hexamer, each loop extending from C-terminal domains of different gp372.

In summary, interactions between MCPs within the same hexamer are mediated mostly by the triangular body, while inter-hexamer interactions involve primarily the hand region of each MCP. Importantly, the hexamers are characterized by a unique 2 nm central pore, whose biological function is unknown.

Each pentamer consists of five MCP monomers (Figure. 2A). In comparison with a gp372 in a hexamer, the N-terminal in pentamer’s gp372 shows a different folding direction (Figure. 2B, left panel). In addition, gp372 in a pentamer changes the relative orientation of the triangular body compared to the MCP in a hexagon (Figure. 2B, right panel).

**Figure 2.**
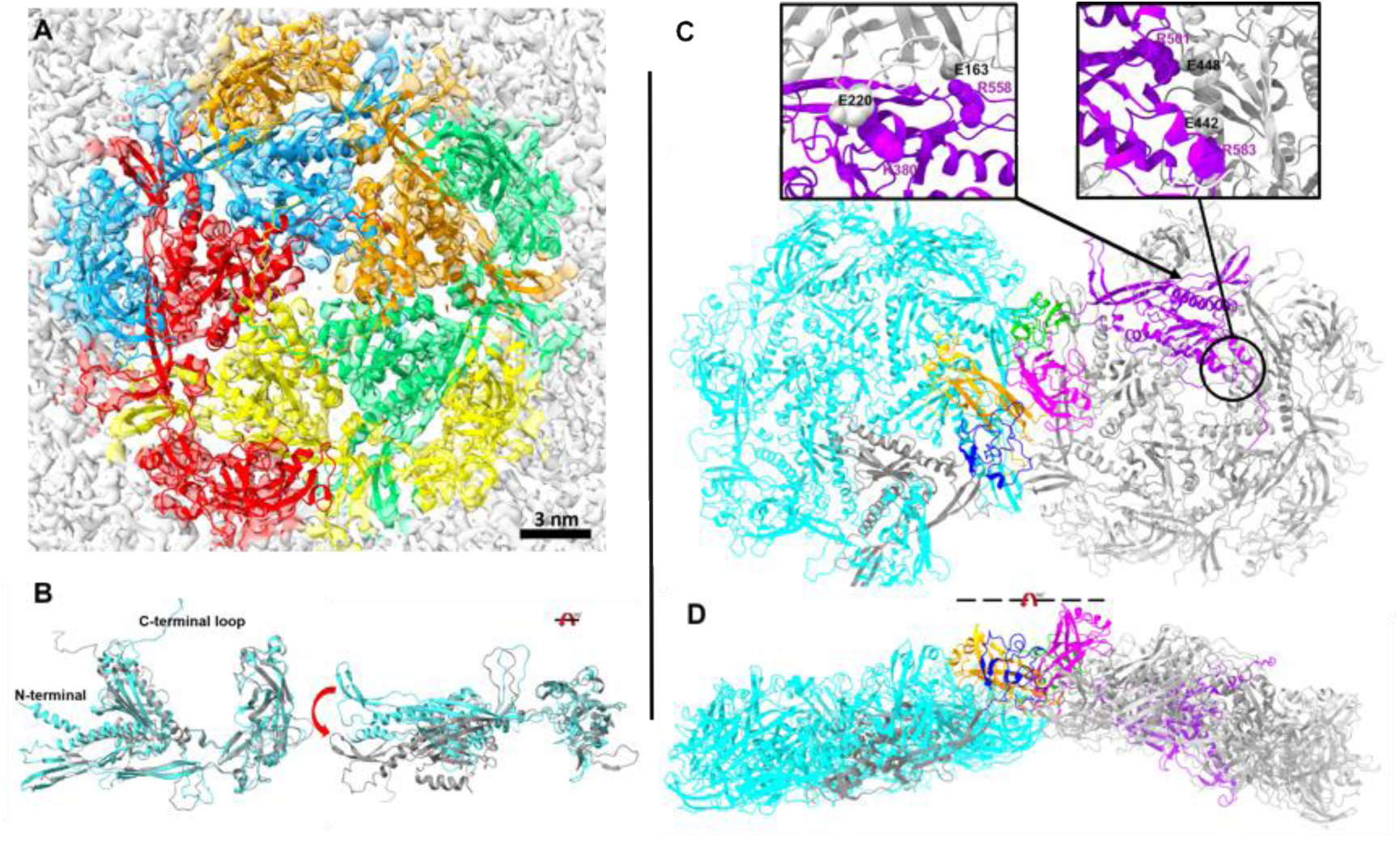
Pentamer arrangement in the capsid of ϕKp24. (A) Top view of a pentameric capsomere. Five gp372 structures were fitted into the density map and colored differently. Compared with the hexamer, the pentamer lacks a structured pore at the center. Each C-terminal loop is flexible, and it tends to move to the area closer to another gp372’s hand. (B) Structural comparison between the MCP (cyan) in a hexamer and MCP (gray) in a pentamer. There are three distinct differences: the C-terminal loop, the N-terminal α helix, and the pendulum angle (red arrow) of two triangular bodies (aligned hand and arm). The right image in B shows the comparison after rotation of the left structure by 90°. (C) Pentamer-hexamer interactions. The interactions between a pentamer (gray) and a neighboring hexamer (cyan) are similar to the interactions between two hexamers. The upper left panel shows that the hand (gray) from the clockwise (CW) neighbor can interact with the triangular body (purple) in the same pentamer using salt bridges (residue E220 and K380, E163 and R558). In the pentamer MCP, residues R501 and R583 in the triangular body interact with residues D448 and E442 (the enlarged rectangular box, upper right panel). The gp372 from a hexamer and the gp372 from a pentamer are highlighted by different colors according to the structural units. The triangular body, arm, and hand of the gp372 in hexamer are colored black, deep blue, and orange. The triangular body, arm, and hand of the gp372 in pentamer are colored purple, green, and pink. (D) Side view of pentamer-hexamer interactions after rotation of the upper image by 90°.

Overall, the interactions of the MCPs in a pentamer are similar to those seen in the hexamers (Figure. 1F). We assembled a list of the putative inter-MCP contacts for the pentamer (Figure. S7). However, as is apparent from the side view, the pentamers are more dome-shaped compared to the flatter arrangement of the hexagons (Figures. 1G and 2D). This increased curvature alters the relative orientation between the triangular body and hand regions in the pentamer MCPs as compared to the hexamer MCP, leading to altered residue-residue contacts. Specifically, in the pentamer MCP, residues R501 and R583 in the triangular body interact with residues D448 and E442, respectively, in the triangular body of the clockwise neighbor. The interactions with the clockwise hand region are reduced overall, involving residues K380 and R558 interacting with residues E220 and D163, respectively. In addition, as seen in the hexamer MCP, the hand region in the pentamer MCP plays an analogous role in mediating hand-hand interactions with a neighboring hexamer through a charged patch of residues described above. In contrast to the hexamer arrangement, the C-terminal loops in the pentamer do not appear to form a well-defined pore in the density map. Instead they are disordered, perhaps owing this different arrangement to the reduced space and greater steric hindrance between the MCPs in this region (Figure 3A). Nevertheless, we do observe a small pore at the center of the pentamer that may also be capable of solvent transport.

**Figure 3.**
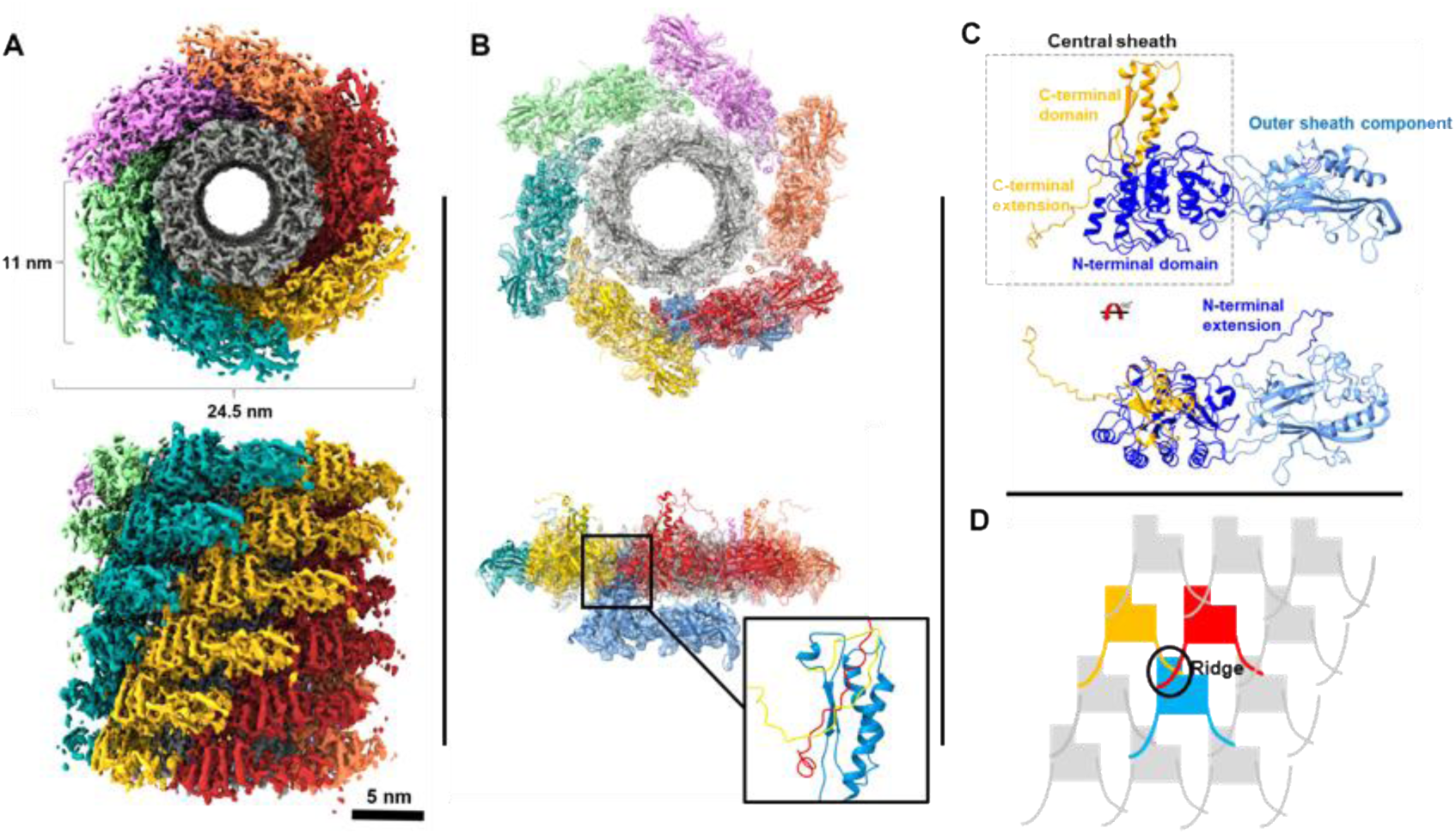
The ϕKp24 helical tail. (A) Top view (top image) and side view (bottom image) orientations of the 3D structure (EMD-14357) of the 4.1 Å helical tail. Each of the six helical strands is colored differently. (B) Model of tail sheath protein gp118 generated from Alphafold2. Sheath protein gp118 and the inner tube protein of phage T4 (PDB ID: 5W5F) were fitted into one ring of the ϕKp24 EM map (upper image, top view; lower image, side view). In the bottom panel, the sheath monomer (blue) that originates from the lower layer is shown to allow visualization of the ring-ring interactions. Contacts between three sheath monomers are highlighted from two successive rings (the enlarged rectangular box, lower right panel). (C) Ribbon diagram of the predicted AlphaFold2 model of the ϕKp24 sheath monomer, gp118. The model consists of two parts: the central sheath (dotted box, orange and dark blue) and an extended outer sheath component (light blue). The lower image is rotated in respect to the upper image by 90° to show the N-terminal extension. (D) Schematic of the ϕKp24 sheath organization. The ridge, where the three sheath monomers from two layers interact, is shown in a circle.

In summary, MCPs interact with each other within pentamers using the triangular body, and with neighboring hexamers using the hand. The structured pore found at the center of hexamers is absent in pentamers due to conformational changes in pentamer’s MCPs.

For the reconstruction of the tail tube and sheath, we picked 400 Å long segments from phages that contained full capsids (Figure. S8B). After 3D refinement, the tail structure (EMD-14357, Figure. 3A) in its extended conformation was determined to a resolution of 4.1 Å in Relion software (Scheres, 2012) (Figure. S8C). The length of the contractile tail is 160 nm on average (measured from the collar to the tip of the spike). The diameter of the tail is 24.5 nm, and the length of the sheath monomer across the longest axis is

11 nm (Figure. 3A). Like other described contractile ejection systems and bacteriophage tails, the contractile tail sheath of ϕKp24 is assembled around the inner tube. The sheath and inner tube follow the same helical symmetry with an additional 6-fold symmetry around the tail axis. The thickness of one hexameric ring is 4 nm, and the rotation between one ring monomer and the next is 30° (Figure. 3A).

Overall, we reconstructed a section of the tail of phage Kp24 in extended conformation at 4.1 Å resolution.

### Structure of sheath protein gp118

The structure of the sheath protein (gp118, accession no. QQV92013.1) was predicted using AlphaFold2 and fitted into the ϕKp24 tail EM map using ISOLDE (Croll, 2018) in ChimeraX (Goddard et al., 2018) (Figure. 3B). The full-length sheath protein is composed of outer and central sheath components. The outer sheath component (Figure. 3C, light blue) is located at the tip of the sheath monomer, pointing outwards. It is a homologue of the bacteriophage phiKZ tail sheath protein gp29PR (PDB ID: 3JOH). This outer sheath component is not involved in the interactions between the subunits during assembly (Aksyuk et al., 2011). In the contracted conformation, the outer sheath components of two layers may interact with each other through electrostatic forces. This is likely since the upward facing surfaces of the outer sheath components contain a negative charged patch, whereas the downward facing surfaces display a positive charged patch (Figure. S9).

The central sheath component (Figure. 3C, orange, and dark blue) consists of the C-terminal domain, N-terminal domain, and two long extensions. The central sheath is homologous to the rod-like (R)-type pyocin sheath protein (PDB ID: 6PYT) and is essential in sheath assembly and contraction. This indicates that the tail of phage ϕKp24 adopts a similar contractile mechanism as the *Pseudomonas aeruginosa* pyocin (Ge et al., 2015). To facilitate the interactions between tail sheath proteins in the tail, the two long extensions from the central sheath (the N-terminal extension, residues M1 – P37; the C-terminal extension, residues F665 – G689) can connect to the C-terminal domains of two adjacent gp118 proteins located in the ring below (Figure. 3D). Additionally, the C-terminal domain can connect with two adjacent sheath proteins located in the upper ring (Figure. 3B, lower right panel). Thus, three sheath proteins from two different layers link together within a so-called ridge (Figure. 3D). The central sheath component cannot interact with each other within a ring in the same layer, and all interactions between rings are confined to the ridges (Figure. 3A, D). The atomic structures show that the sheath extensions can create a “mesh” (Figure. 3D). The sheath proteins likely remain linked with each other during the transition from an extended to a contracted tail similar to the mechanism of pyocin contraction (Ge *et al*., 2015).

Overall, the tail sheath protein is composed of outer and central sheath components. The central sheath component is essential in sheath assembly and contraction.

The tube of phage ϕKp24 is homologous to the tube of other contractile injection systems, such as phage T4 (Zheng et al., 2017b), R-type pyocins (Ge *et al*., 2015), *Serratia entomophila* antifeeding prophage (afp) (Heymann et al., 2013), and the bacterial Type VI Secretion System (T6SS) (Clemens et al., 2015; Kudryashev et al., 2015). These different tubes show the ability to translocate different substrates and the nature of the substrate determines the properties of the tube’s channel (Zheng *et al*., 2017b).

Here we used the phage T4’s tube protein monomer (Figure. 4A, PDB: 5W5F) for model building. The model was then flexible fitted into the tube EM map using ISOLDE in ChimeraX. One ring-like structure consists of six tube proteins, which represents an assembly unit of the inner tube of phage ϕKp24 (Figures. 4B and 4C). The ring structure shows a complementary surface charge on its contact interface: one is positively charged, and the other is negatively charged. This can create an electrostatic dipole which is related to the directional self-assembly (Ge *et al*., 2015). The inner surface of the tube displays a prominent negative charge (Figure. 4D), which prevents the negatively charged phage DNA sticks to the surface.

**Figure 4.**
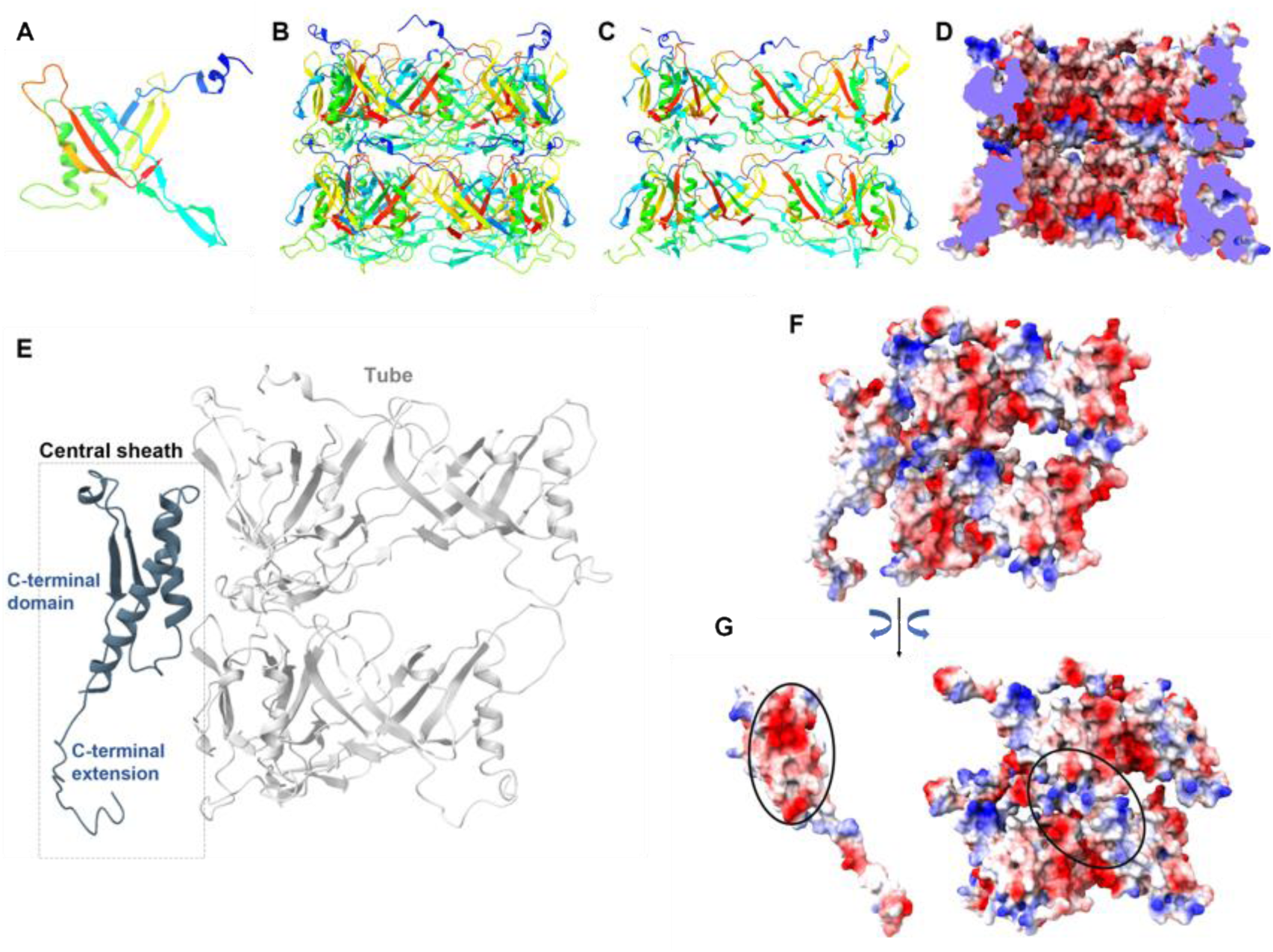
Inner tube structure of phage ϕKp24 and sheath-tube interaction. (A) Ribbon diagram of T4’s tube protein monomer (PDB 5W5F) for model building. (B) Ribbon diagram of two discs (hexamers) in the tube of phage ϕKp24. (C) Cut-away view as ribbon diagram showing the inner surface of the tube structure in B. (D) Cut-away view of electrostatic diagram showing the inner surface of the tube structure in B. Charge distribution: red = negative; blue = positive; white = neutral. Electrostatic potential was colored in ChimeraX. (E) Side views of the interface between a part of sheath protein (C-terminal domain of central sheath) and four tube-protein subunits. (F) The charged surface of E. (G) An open-book view of F. The complementary patches of interacting charges on both sheath part and tube are marked with ovals.

In summary, the structure and function of the tube of phage ϕKp24 are similar to that of the tube in phage T4.

### Sheath-tube interaction

Sheath–tube interactions may arise from electrostatic and non-covalent forces and from viscosity in the nanoscale gap (interstitial water) between the tail tube and the surrounding sheath (Maghsoodi et al., 2019). The electrostatic forces are largely perpendicular to the tail tube axis and thus contribute to the connection between the sheath and tail tube. Additionally, the sheath protein connects with inner tube subunits using electrostatic interaction (Figure. 4G). The outer surface of the tube subunits displays a positive charged patch (Figure. 4G, right panel). The C-terminal domain of the sheath protein binds to the tube’s patch via a complementary negative-charged patch on one of its two α-helices.

### The model for tail assembly

The tube is known to have a critical role in the assembly of the sheath in the extended state of phage tails (Leiman and Shneider, 2012). The assembly of ϕKp24’s sheath starts from the baseplate, as is the case for phage T4 (King and Mykolajewyoz, 1973). The tube and the baseplate can form a “platform” to which the first disc of sheath subunits bind. Subsequently, the tube and the disc of extended sheath subunits serve as a scaffold for assembly of the rest of the sheath. Thus, the assembly of the contracted state is avoided by the creation of a template in which sheath subunits interact along the ridge without any lateral contacts (Figure. 3D). This template-driven assembly probably results in a metastable oligomeric structure, which could be unlocked by the baseplate for tail contraction upon interaction with the target-cell surface.

### Tomographic reconstruction and segmentation of tail fibers

To investigate the structures and the conformational changes of the tail fibers during the cell attachment and genome injection, we used cryo-ET. We imaged ϕKp24 together with its *K. pneumoniae* host and observed intact phages in four different states: free particles with and without DNA, and adsorbed particles with and without DNA. These four states correspond to four different states during phage infection: free phage, early-infection phages attached to the host, post-infection (empty) phages still attached to the cell, and post-infection (empty) free phages. A representative 2D cryo-ET image (Figure. 5A) shows several of these states. To find the architectures of disordered fibers at different moments during the process of infection, we focused on intact ϕKp24 attached to a *K. pneumoniae* cell. We selected and extracted intact phages from 89 denoised 3D reconstructions in IMOD (Kremer et al., 1996).

**Figure 5.**
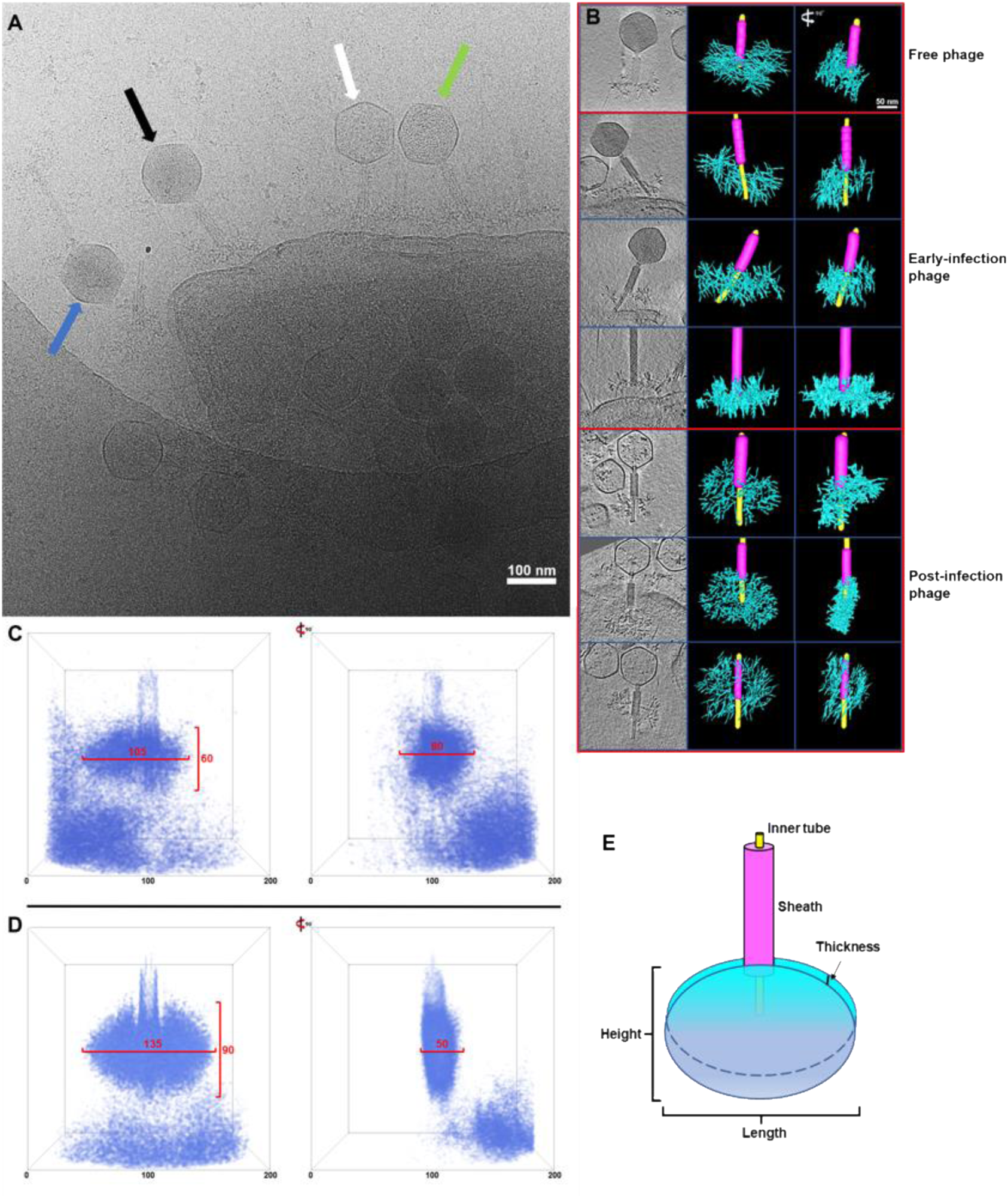
Cryo-ET of phage ϕKp24 infecting *K. pneumoniae* and averaged 3D structures of tail fibers. (A) 2D cryo-ET image showing several different states of phage ϕKp24 in its native environment: free phages (phages with dark capsids, blue arrow), early-infection phages (phages with dark capsids and attached to host membrane, black arrow), mid-infection phages (phages with half-dark capsids and attached to host membrane, green arrow), and post-infection (empty) phages still attached to the cell (white arrow). (B) Tomograms of ϕKp24 and manual segmentations. The first column shows seven representative tomograms of ϕKp24 in different states during the injection. From top to bottom, the first tomogram shows a free phage (full capsid), the second to fourth show early-infection phages (full capsids) attached to a cell, and the fifth to seventh show post-infection phages (empty capsids) attached to the host. The second column shows the manual segmentations of the tail fibers corresponding to the tomograms shown on the left. The sheath is colored in purple, fibers are colored in cyan, and the tube is colored in yellow. The third column shows the side view of every left segmentation after rotation by 90°. (C) 3D superimposed structure of tail fibers in pre-infection phages. 89 tail-fibers structures of pre-infection phages were aligned and averaged. The right image shows the side view of the structure after rotation by 90°. Both the outer box and red line are scale bars with pixel unit. The outer square box size is 200 pixels, pixel size is 1.312 nm. The size of the tail-fiber structure is about 137.8 nm in length, 78.7 nm in height, and 104.9 nm in thickness. (D) 3D superimposed structure of tail fibers in post-infection phages. 338 tail fiber structures of post-infection phages attached to host were aligned and averaged. The tail fiber structure resembles a flattened disc. The size is 177.1 nm in length, 118.1 nm in height, and 65.6 nm in thickness. (E) Schematic diagram of phage sheath (purple), inner tube (yellow), and structure of tail fibers (cyan). Here we defined the height, length, and thickness of the structure.

Because of the high complexity and heterogeneous nature of the tail fibers, we chose to manually segment the tail fibers using the IMOD segmentation function (Kremer *et al*., 1996). We first extracted sub-volumes of individual phages from the whole tomograms (Figure. 5B). Here, we aimed at selecting phages that were in different stages of infection, from the pre- to the post-infection state. We then segmented the phage tails of these phages (Figure. 5B) and compared their overall architecture. We found that fibers of pre-infection phage are arranged around the baseplate in a near-spherical organization. In contrast, when phages attach to a host cell, the fibers interact with binding sites on the cell envelope, changing the overall architecture. The tail fibers arrange along the cell surface, creating a flat “tail plate”. In addition, we observe that the tail of the attached phages is not vertically arranged with respect to the cell envelope. Instead, it is tilted, while this could be an artifact of sample preparation, it is also possible that the injection of the genome occurs at an angle in some cases.

### Tail fiber analysis using machine learning

Due to the time-consuming manual segmentation of the tail fibers, we built a neural network to auto-detect the tail fibers of phage ϕKp24 in reconstructed tomograms. We trained the network using the manual segmentations of the tail fibers and, after several rounds of optimization, applied the trained network to all tomograms, generating 3D maps containing tail fiber structures. After extraction and format conversion, 89 tail fiber structures of pre-infection phage and 338 tail fiber structures of post-infection attached phage were generated. 40 representative tail fiber structures of pre-infection and tail fiber structures of post-infection are shown in Figures. S10 and S11. These 3D structures show clearly that each tail fiber structure is in a different conformation. In the pre-infection phage group (Figure. S10), the tail fibers are arranged around the baseplate in a near-spherical arrangement. The size range of the tail fiber structures is approximately 110-130 nm in height, 150-170 nm in length, and 90-130 nm in thickness. In the post-injection phage group (Figure. S11), most of the tail fiber structures arrange in an oval, disc-like configuration. The size range in 3D is 120-150 nm in height, 160-190 nm in length, and 60-100 nm in thickness.

To further reveal the conformational changes of tail fibers in pre-infection and post-infection stages, the maps within each group were aligned to each other, and averaged to produce ‘canonical’ fiber shapes. Finally, we got the 3D superimposed structures of tail fibers in pre-infection (Figure. 5C) and post-infection (Figure. 5D), which clearly show a change in tail fiber conformation from a near-spherical shape to a flat disc.

Between these configurations, the thickness changes by approximately 40 nm. In addition, the tail from the post-infection superimposed structure is not perpendicular to the fibers’ disc, which suggests that the tail is not oriented perpendicular to the host surface during DNA injection. This is also apparent in the individual structures of tail fibers and manual segmentations.

Overall, manual and machine learning segmentation of the tail fibers revealed a significant conformational change of the tail fiber complex upon binding to the host cell surface, likely to position the phage for DNA injection.

A collection of strains with 77 different capsular (K-type) serotypes was used to characterize the infectivity of ϕKp24. This analysis revealed nine serotypes (K2, K13, K19, K25, K35, K46, K61, K64, and K81) susceptible to ϕKp24 with an efficiency of plating ranging from 1 - 0.001 compared to the ϕKp24 host strain (Figures. S12 and S13). However, sequence similarity (Altschul et al., 1990) and structure-based homology modeling (Kelley et al., 2015) suggest that the phage carries 14 tail fiber proteins (Table 1, gp168, gp196, gp294, gp295, gp300, gp301, gp303, gp304, gp306, gp307, gp308, gp309, gp310, and gp313) instead of the nine previously predicted ones (Bonilla *et al*., 2021). These proteins possess putative depolymerizing activity (lyase or hydrolase) and a β-helical fold, which is a hallmark feature for *Klebsiella* phage depolymerases. Modeling of the 14 proteins with RoseTTAFold (Baek et al., 2021) revealed a typical elongated protruding shape composed of parallel β-strands orthogonal to the long axis (Figure. 6). For five of the depolymerases, a putative K-serotype can be predicted based on amino acid sequence similarity to experimentally verified depolymerases (Table 1). Despite the low level of identity (below 51%) we can assume the identity is not incidental, since phage ϕKp24 was also able to infect *Klebsiella* spp. with the corresponding capsular serotypes (Table 1).

**Figure 6.**
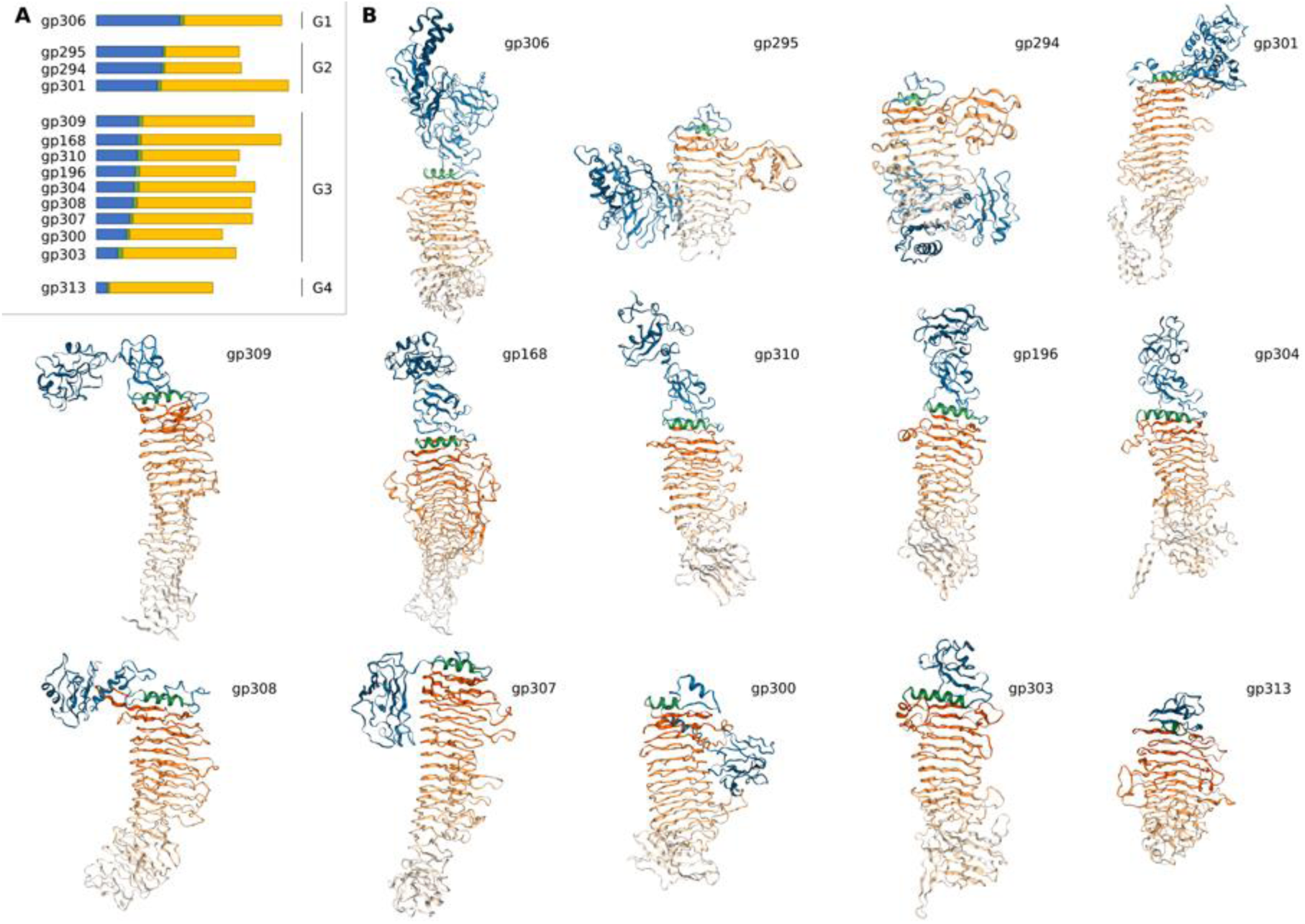
*In silico* structural analyses of tail fibers with predicted depolymerase activity. (A) Schematic delineation of the different modules in fourteen predicted tail fibers results in four groups (G1, G2, G3, and G4) according to the length of their structural modules. Note that the relative orientations of the individual domains in the predicted structure can be different than depicted. (B) 14 predicted tail fiber structures. Tail fibers with depolymerase activity are typically composed of structural modules responsible for tail attachment and branching (blue), a separating α-helix (green), and a β-helix with enzymatic activity (yellow).

**Table 1.**
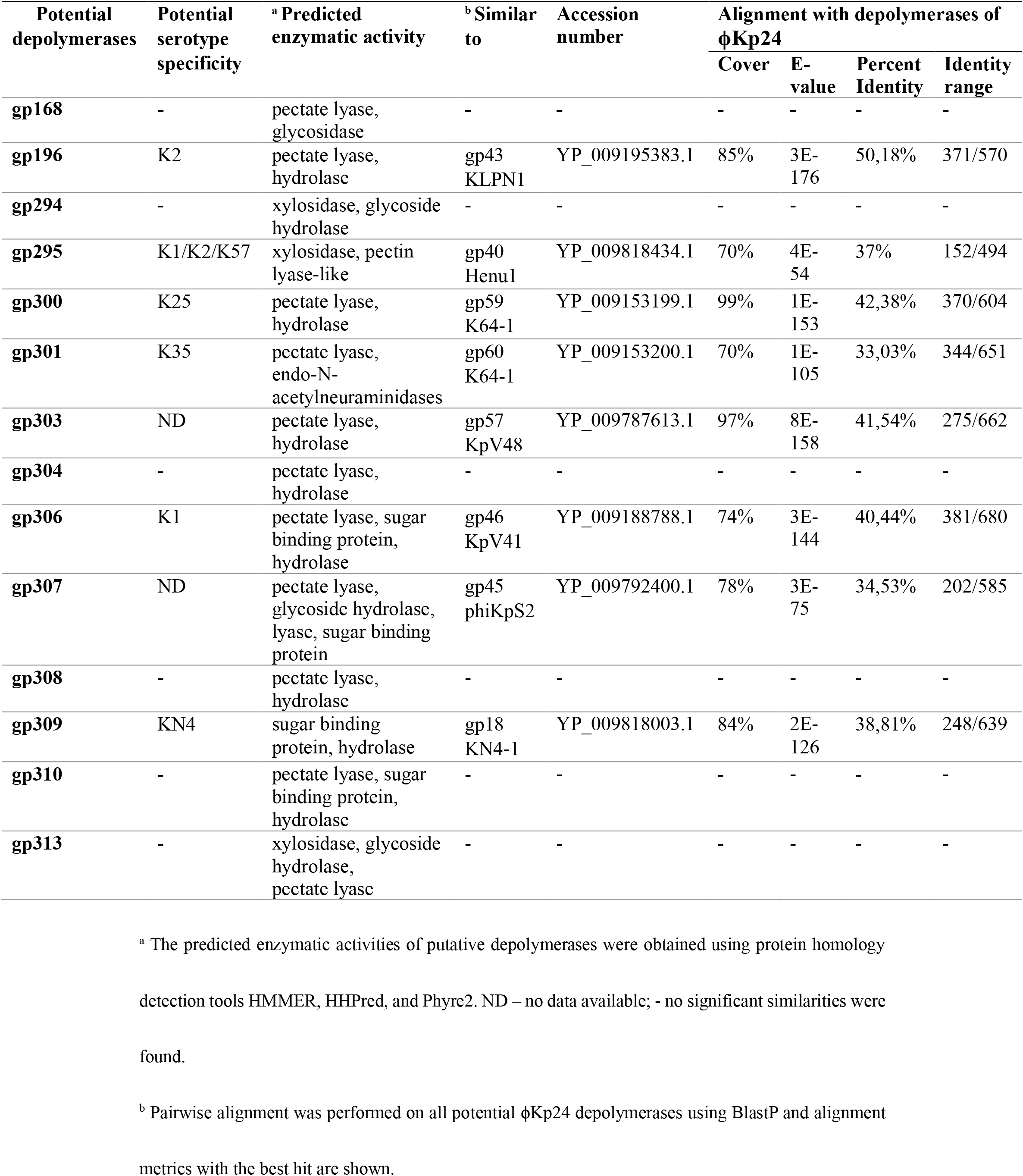
Predicted K-serotype specificity and enzymatic activity of putative depolymerases.

| Potential depolymerases | Potential serotype specificity | <sup>a</sup> Predicted enzymatic activity | <sup>b</sup> Similar to | Accession number | Alignment with depolymerases of ϕKp24 | | | |
| --- | --- | --- | --- | --- | --- | --- | --- | --- |
|  |  |  |  |  | Cover | E-value | Percent Identity | Identity range |
| **gp168** | - | pectate lyase, glycosidase | - | - | - | - | - | - |
| **gp196** | K2 | pectate lyase, hydrolase | gp43 KLPN1 | YP_009195383.1 | 85% | 3E-176 | 50,18% | 371/570 |
| **gp294** | - | xylosidase, glycoside hydrolase | - | - | - | - | - | - |
| **gp295** | K1/K2/K57 | xylosidase, pectin lyase-like | gp40 Henu1 | YP_009818434.1 | 70% | 4E-54 | 37% | 152/494 |
| **gp300** | K25 | pectate lyase, hydrolase | gp59 K64-1 | YP_009153199.1 | 99% | 1E-153 | 42,38% | 370/604 |
| **gp301** | K35 | pectate lyase, endo-N-acetylneuraminidases | gp60 K64-1 | YP_009153200.1 | 70% | 1E-105 | 33,03% | 344/651 |
| **gp303** | ND | pectate lyase, hydrolase | gp57 KpV48 | YP_009787613.1 | 97% | 8E-158 | 41,54% | 275/662 |
| **gp304** | - | pectate lyase, hydrolase | - | - | - | - | - | - |
| **gp306** | K1 | pectate lyase, sugar binding protein, hydrolase | gp46 KpV41 | YP_009188788.1 | 74% | 3E-144 | 40,44% | 381/680 |
| **gp307** | ND | pectate lyase, glycoside hydrolase, lyase, sugar binding protein | gp45 phiKpS2 | YP_009792400.1 | 78% | 3E-75 | 34,53% | 202/585 |
| **gp308** | - | pectate lyase, hydrolase | - | - | - | - | - | - |
| **gp309** | KN4 | sugar binding protein, hydrolase | gp18 KN4-1 | YP_009818003.1 | 84% | 2E-126 | 38,81% | 248/639 |
| **gp310** | - | pectate lyase, sugar binding protein, hydrolase | - | - | - | - | - | - |
| **gp313** | - | xylosidase, glycoside hydrolase, pectate lyase | - | - | - | - | - | - |

<sup>a</sup> The predicted enzymatic activities of putative depolymerases were obtained using protein homology detection tools HMMER, HHpred, and Phyre2. ND – no data available; - no significant similarities were found.
<sup>b</sup> Pairwise alignment was performed on all potential ϕKp24 depolymerases using BlastP and alignment metrics with the best hit are shown.

The N-terminus of *Klebsiella* phage tail fibers typically has a structural role with domains involved in either attachment to the phage tail or another tail fiber or providing a branching site for other tail fibers (Latka *et al*., 2019). The C-terminus contains the enzymatic depolymerase domain defining capsule serotype specificity and features a β-helical fold. The C-terminus may also include additional chaperone or carbohydrate binding domains (Cornelissen et al., 2011; Latka et al., 2017; Schwarzer et al., 2012; Weigele et al., 2003; Yan et al., 2014). The N- and C-terminus are typically separated by an α-helix (Seul et al., 2014) (Figure. 6). The 14 putative depolymerases were divided into four groups based on the length of the N-terminal structural part. The rationale behind this grouping is that a long N-terminal structural part is indicative for a role as primary tail fiber (i.e., the tail fiber attaching the full tail fiber system to the phage tail) including multiple domains involved in direct attachment to the phage tail and providing additional branching sites for secondary tail fibers. These secondary tail fibers dock with shorter N-terminal structural domains as described for the branched tail fiber systems of phage G7C (Prokhorov et al., 2017) and CBA120 (Plattner et al., 2019). Secondary tail fibers can comprise additional docking sites for further branching to tertiary tail fibers as described for phages belonging to the Menlow group (Latka *et al*., 2019). As such, the tail fibers of ϕKp24 can be clustered into four different groups corresponding to more peripheral positions in the whole tail fiber architecture, starting from the central phage tail. Gp306 is predicted as the primary tail fiber (group 1) directly connecting to the phage tail with its 398 aa-long N-terminus. HHPred (Zimmermann et al., 2018) analysis detected the presence of a T4gp10-like fold in the region of 61-275 aa of the gp306 N-terminus (Table S1). T4gp10-like folds are reported to function as branching sites for additional tail fibers in phage G7C and CBA120. However, similarities are generally low and mainly at the tertiary structure level, rendering them less amenable for prediction at such branching sites. The long T4gp10- like fold may provide branching sites for more than one tail fiber from group 2 (gp294, gp295 and gp301 with 316, 311 and 289 aa-long N-termini, respectively). At least in one of these (gp301), a T4gp10-like fold could be predicted (Table S1) but the similar length of the N-termini of gp294 and gp295 suggests the presence of branching sites as well. Group 3 contains nine proteins (gp168, gp196, gp300, gp303, gp304, gp307, gp308, gp309 and gp310) with lengths of the N-terminal structural parts ranging from 99 aa (gp303) to 199 aa (gp309), being sufficient to harbor a domain responsible for attachment to a branching site. Only one tail fiber with predicted depolymerase activity possesses a short peptide (51 aa) preceding the separating α-helix, i.e., gp313 (group 4). This N-terminal peptide may be sufficiently long to mediate attachment to another tail fiber; however, it is also possible that gp313 is not integrated in the phage particle but occurs only in a soluble, non-integrated form. Altogether, the tail fiber architecture of ϕKp24 appears to be organized in different peripheral layers with gp306 acting as the primary tail fiber anchoring the whole tail fiber system to the tail.

## DISCUSSION

The increasing problem of antibiotic resistance in bacterial pathogens generated renewed interest in the use of phages to treat antibiotic-resistant bacterial infections. For the optimal use of phages in therapeutic applications, we need to improve our understanding of their structure and function. The recently described *Klebsiella* phage ϕKp24 is a promising candidate phage with an expanded host range, infecting at least nine different capsular types and likely more based on the fourteen predicted tail fibers with depolymerase activity. However, unraveling its unique morphology represents a challenge for detailed structural studies. Especially the large size of the capsid and the heterogeneous nature of the tail fibers pose a challenge for data collection and analysis. They require the combination of different structural methods to solve the overall structure. Protein structure prediction algorithms, such as AlphaFold2 and RoseTTAFold have become invaluable tools to cryo-EM based macromolecular structure elucidation. Furthermore, machine learning provides a new way to analyze intricate image data of cryo-ET. In this study, we combined single particle analysis and cryo-electron tomography with protein structure prediction, molecular simulations, and machine learning. This allowed us to uncover the structure of ϕKp24 particle, as well as characteristic morphological changes the phage undergoes during host infection.

The large size of especially the capsid together with a large amount of data also pose computational challenges during data processing. More specifically, the diameter of ϕKp24’s capsid is around 1450 Å. According to the falling gradient of the FSC curve in Relion, the dataset used for final 3D refinement had to be binned by 1.5x to balance the final resolution with the available computational resources. The capsid of ϕKp24 is currently the highest resolution capsid of any jumbo phage deposited in the EMDB. The main structural component of the capsid is the MCP gp372. This unusually large protein is the basis for the hexamers and pentamers that form the capsid. Typically, phage capsids are composed of one (or two) major capsid proteins (MCPs) and decoration proteins (additional external proteins that link the MCPs together). Our reconstruction of ϕKp24’s capsid has shown that the entire capsid of ϕKp24 can be assembled by a single MCP alone.

In the hexamers, the six long C-terminal loops form a pore in the center of a hexamer. To our knowledge, a similar pore has not been reported before in other published jumbo phage capsids. The exact function of this pore is currently unclear. We speculate that the pore may be involved in balancing pressure differences during loading of the DNA into the capsid, or during DNA ejection when capsid size increases.

There is an evolutionary link emerging between some phage tail proteins and the Gram-negative bacterial type VI secretion system (T6SS) which is implicated in various virulence-related processes (Basler, 2015; Leiman et al., 2009; Records, 2011). The structure of the contractile sheath reported here shows strong similarities to other contractile systems, such as the R-type pyocin (Ge *et al*., 2015) and T4 (Zheng *et al*., 2017b). This suggests that the helical tail of phage ϕKp24 functions in a similar fashion. Furthermore, the structure solved here suggests that the subunit-connecting extensions (C terminal extension) of the sheath protein gp118 function as hinges during contraction, with the central sheath of gp118 likely maintaining its structure. The rearrangement of sheath subunits upon contraction leads to a much more densely packed structure. Here, the ridges are brought closer together to become partly interdigitated. In the contracted tail, the connections with the inner tube are lost, allowing the tube to insert into the target cell membrane during infection (Ge *et al*., 2015).

The tomograms reveal that the tails of attached phages do not appear vertical in respect to the cell envelope. Instead, the tails of phages with full or empty capsids are tilted. While we cannot rule out that this observation is an artifact from the sample preparation, this attachment may be physiologically relevant. A tilted tail may reduce the required energy for contraction and DNA ejection, as well as the damage of the cell envelope to prevent premature lysis of the host.

Unraveling the structure of the tail fibers in ϕKp24 was especially challenging. Unlike the capsid and tail, the tail fibers are structurally highly heterogeneous and not amenable for single-particle analysis. By using a neural network that is specifically designed for accurate training with a limited number of examples (Pelt and Sethian, 2018), we were able to train a network using segmentations of the tail fibers of only seven phage particles, thereby limiting labor-intensive manual segmentation. After training, the network enabled automatic analysis of the tail fibers of more than 600 phages, demonstrating the possibilities of machine learning for aiding the analysis of complex structures.

The tail fiber data revealed the exquisite complexity of these tail fibers and a significant change in their arrangement between free phages and those bound to a host cell. While the fibers arrange as a sphere around the tail tip of a free phage, in the host-bound phages the tail fibers adjust to the host envelope and form a plate. Notably, the change in the conformation of the tail fibers precedes DNA release. Therefore, it may be part of the triggering mechanism itself.

Host range analysis of ϕKp24 revealed an unprecedented number of capsule serotypes that are host for this phage. This observation corresponds to the multiple tail fibers with specific depolymerase activity predicted in the large phage genome and the highly branched tail fiber system that could be visualized. Structural analysis of the tail fibers indicates an extensive structural architecture to integrate all tail fibers in the phage particle with gp306 emerging as the primary tail fiber candidate making direct contact with the phage tail, while subsequent groups of tail fibers have an increasingly shortened structural domain, hinting at a more peripheral location.

## ACKNOWLEDGMENTS

This work is part of the research program National Roadmap for Large-Scale Research Infrastructure 2017–2018 with project number 184.034.014, which is financed in part by the Dutch Research Council (NWO). CKC and PJS are funded by BBSRC, MRC, and Wellcome. The depolymerase experiments and analyses were supported by National Science Centre, Poland, with grant numbers UMO-2017/26/M/NZ1/00233 to ZD-K, AL, and UMO-2020/38/E/NZ8/00432 to AO. AL holds a junior postdoctoral fellowship of the Research Foundation – Flanders (FWO) with grant number 1240021N. DMP is supported by NWO with grant number 016.Veni.192.235. SJJB is supported by the NWO with VICI grant VI.C.182.027 and the European Research Council (ERC) CoG with grant no. 101003229. RO was supported by the China Scholarship Council (CSC) with project number 201906280465.

We thank Jamie Depelteau and Adam Sidi Mabrouk for cryo-EM sample preparation help; Wen Yang and Willem Noteborn for cryo-EM data collection; Ludovic Renault and Frédéric Bonnet for cryo-EM data processing help; Boris Estrada-Bonilla for technical support; Xiao Zhang for help with extraction of the network results and Lei Zhang for critical feedback on the manuscript.

The *K.* pneumoniae reference strains used in this study (CIP labelled strains) were obtained from the Collection de l’Institut Pasteur (CIP, Paris, France).

## SUPPLEMENTAL INFORMATION

### Phage infection movie

https://www.dropbox.com/s/26l8m0f5a3rlf26/Flowerphage%20infection_v005.mp4?dl=0

### Capsid structure movie

https://www.dropbox.com/s/hh9z5ldskp9xgkv/Capsid%20structure%20organization-v039.mp4?dl=0

### Supplemental material

https://www.dropbox.com/s/c3fn2zxuhpbj7fj/Flower%20phage%20supplementary%20V11.pdf?dl=0

## AUTHOR CONTRIBUTIONS

Conceptualization A.B., S.J.J.B, Y.B., Z.D.K.; Methodology A.B., R.O., Y.B., Z.D.K., C.K.C.; Software R.O., C.K.C; Formal analysis R.O., C.K.C, D.M.P., A.L., A.O, V.W.; Investigation R.O., A.R.C., C.K.C., A.O.; Resources A.B., S.J.J.B, D.M.P, Y.B., Z.D.K.; Data curation R.O., A.B., Y.B., Z.D.K.; Writing original draft R.O., A.B., Y.B., D.M.P., Z.D.K.; Writing- review and editing R.O., A.R.C., C.K.C., V.W., P.J.S., D.M.P., S.J.J., Y.B., A.L., Z.D.K., A.O.; Visualization R.O., V.W., Y.B.; Supervision A.B., S.J.J.B, Y.B., .; Project Administration A.B.; Funding acquisition A.B., S.J.J.B., C.K.C., P.J.S., Z.D.K., A.L., D.M.P., O.R.

## RESOURCE AVAILABILITY

### Lead contact

Further information and requests for resources and reagents should be directed to and will be fulfilled by the lead contact, Ariane Briegel.

### Materials availability

This study did not generate new unique reagents.

### Data and code availability

• Cryo-EM reconstructed data have been deposited at EMDB (the Electron Microscopy Data Bank) and are publicly available as of the date of publication. Accession numbers are listed in the key resources table. Microscopy, Protein prediction data reported in this paper will be shared by the lead contact upon request.
• All original code has been deposited at GitHub and is publicly available as of the date of publication. DOIs are listed in the key resources table.
• Any additional information required to reanalyze the data reported in this paper is available from the lead contact upon request.

## EXPERIMENTAL MODELAND SUBJECT DETAILS

### Bacterial strains

The *K. pneumoniae* reference strains used in this study (Figure S12) were obtained from the Collection de l’Institut Pasteur (CIP, Paris, France, CIP labelled strains) or purchased from The National Collection of Type Cultures (NCTC labelled strains). Bacteria were stored at −70 °C in Trypticase Soy Broth (TSB, Becton Dickinson and Company, Cockeysville, MD, USA) supplemented with 20% glycerol.

### Bacteriophage

Phage ϕKp24 (Genbank accession no. MW394391) was previously isolated from sewage water (Bonilla *et al*., 2021). ϕKp24 was cultured in 2 L of Lysogeny Broth (LB) using *K. pneumoniae* strain K5962 as the host (Genbank Bioproject accession no. PRJNA745534). The resulting culture was centrifuged (9,000 × g, 15 min) and the phage-containing supernatant was filter-sterilized. The phage lysate was then washed with SM buffer (100 mM NaCl, 8 mM MgSO4, 50 mM Tris-HCl pH 7.5) and 10× concentrated using a tangential flow cassette (100 kDa PES Vivaflow 200, Sartorius, Germany). Phage titer was determined by spotting serial dilutions of the phage solution on double layer agar (DLA) plates of the K5962 strain for the detection of phages.

## METHOD DETAILS

### Phage K-serotype specificity

The spot method was used to determine the serotype specificity and potential production of capsule targeting depolymerases by phage ϕKp24. The analyses were performed on the *K. pneumoniae* serotype collection (Figure. S12 and S13). The overnight bacterial cultures were suspended in fresh Tryptic Soy Broth (TSB), incubated for 2 hours at 37 °C with agitation (140 rpm) and were poured onto Trypticase Soy Agar (TSA, Becton Dickinson and Company, Cockeysville, MD, USA) plates. After drying, 10 µL of ten-fold serial dilutions of phage ϕKp24 (10^9^ pfu/mL as starting concentration) were spotted on bacterial lawns. After overnight incubation at 37 °C, lawns were inspected for plaque and halo zones. The efficiency of plating was calculated by dividing the titer of the phage at the terminal dilution on the test strain by the titer of the same phage on its host strain.

### Bioinformatic analysis and structural modeling of putative tail fiber genes

The genomic sequence of the phage ϕKp24 was obtained from the GenBank database (NCBI Accession No MW394391). All proteins of phage ϕKp24 were analyzed with Phyre2 (Kelley *et al*., 2015) in search of putative tail fibers with depolymerase activity. Criteria for the prediction of putative depolymerase activity were as in (Latka *et al*., 2019) with some modifications: (1) the protein is longer than 200 residues; (2) the protein is annotated as tail/tail fiber/tailspike/hypothetical protein in the NCBI database; (3) the protein shows homology to domains annotated as lyase [hyaluronate lyases (hyaluronidases), pectin/pectate lyases, alginate lyases, K5 lyases] or hydrolase (sialidases, rhamnosidases, levanases, dextranases, xylanases, glucosidase, galacturonase, galacturonosidase, glucanase) with a confidence of at least 40% in Phyre2; (4) the length of homology with one of these enzymatic domains should span at least 100 residues; (5) a typical β-helical structure should be predicted by Phyre2. Predicted proteins were modeled with RoseTTAFold (Baek *et al*., 2021). The predicted models were further analyzed and the last α-helix before the β-helical structure was chosen as an ending point for the N-terminal structural part. Proteins were ranked by the length of the N-terminal structural part, starting with the longest one.

The proteins were further analyzed with BlastP (Altschul *et al*., 1990) and HMMER (Finn et al., 2011). BlastP was used to determine the serotype specificity based on the amino acid sequence similarities of the depolymerases found in the database. The following criteria were determined for the analysis: (i) the homologous protein should have a confirmed depolymerase activity and specificity for the particular *K. pneumoniae* serotype, (ii) the amino acid sequence identity should exceed 30%, and (iii) the length of the homology should be more than 100 amino acids. The N-terminus of each selected protein was analyzed with HHPred (Gabler et al., 2020; Zimmermann *et al*., 2018) using pairwise alignment in search of T4gp10-like domains. Default parameters were used: MSA generation method: HHblits=>UniRef30; MSA generation iterations: 3; E-value cutoff for MSA generation: 1e-3; Min seq identity of MSA hits with query (%): 0; Min coverage of MSA hits (%): 20; Secondary structure scoring: during_alignment; Alignment Mode:Realign with MAC: local:norealign; MAC realignment threshold: 0.3; Max target hits: 250; Min probability in hitlist (%): 20. The alignment was performed against T4gp10 (NP_049768.1), the N-terminus of CBA120gp163 (orf 211, YP_004957865.1), CBA120gp165 (orf 213, YP_004957867.1) and G7Cgp66 (YP_004782196.1).

### Sample preparation for cryo-ET

*K. pneumoniae* K5962 was grown overnight at 30 ℃ with shaking at 180 rpm. The overnight culture was used to inoculate 10 ml of fresh LB media and grown for 2 hours at 30 °C, 180 rpm. Subsequently, 1.6 ml of phage ϕKp24 (9x10^12^ pfu/ml) were added to the culture and incubated for an additional 80 min. The culture was visually checked to confirm cell lysis, and 200 µl were centrifuged (5000 × *g*, 5 min) and the pellet was re-suspended in the same volume of fresh media. Finally, 5 µl of 10 nm-sized gold beads (Cell Microscopy Core, Utrecht University, Utrecht, The Netherlands) were added to 50 µl of the prepared bacterial-phage mixture. Using the Leica EM GP (Leica Microsystems, Wetzlar, Germany), 3.8 µl of the mixture were applied to a glow discharged Quantifoil R2/2, 200 mesh Cu grid (Quantifoil Micro Tools GmbH, Jena, Germany) and incubated 30 s before blotting at 20 °C with approximately 95% relative humidity. The bacterial-phage grids were blotted for 1s and automatically plunged into liquid ethane. Vitrified samples were transferred to storage boxes and stored in liquid nitrogen until use.

### Sample preparation for SPA

ϕKp24 was concentrated as follows: 1000 µl phage ϕKp24 liquid (9×10^12^ pfu/ml) was spun down at a lower speed (5 k) for 2-3 times (15 min/per spin) until buffer stopped eluting from a protein concentrator (10 kDa cellulose filter). Subsequently, the sample was immediately used for plunge freezing. Using the Leica EM GP, 3.5 µl of the concentrated phage was added to a glow discharged Quantifoil R2/2, 200 mesh Cu grid, incubated 10 s at 18 °C with approximately 95% relative humidity. The phage grids were blotted for 0.7 s and automatically plunged into liquid ethane. Vitrified samples were stored in liquid nitrogen until use.

### Imaging conditions of Cryo-ET

The grids containing *K. pneumoniae* K5962 and ϕKp24 were clipped and loaded into a Titan Krios (Thermo Fisher Scientific (TFS)) equipped with a K3 BioQuantum direct electron detector and an energy filter (Gatan, Inc), which was set to zero loss imaging with a slit width of 20 eV. Imaging targets were chosen by selecting bacterial cells that were in a hole of the carbon film of the EM grid. Bacteria that appeared less dense at low magnification indicated progressed phage infection. Data were collected as movie frame stacks using SerialEM set to a dose symmetric tilt scheme between −54° and 54°, with 2° tilt increments (Hagen et al., 2017; Mastronarde, 2005). The selected nominal magnification was 26,000, which corresponds to a pixel size of 3.28 Å in the collected images. The defocus of the collected data ranged from −4 to −6 µm. The estimated total dose per tilt series was 100 e/ Å ^2^.

### Imaging conditions of SPA

ϕKp24-containing grids were clipped and loaded into a Titan Krios electron microscope (Thermo Fisher Scientific, TFS) operated at 300kV, equipped with a Gatan K3 BioQuantum direct electron detector (Gatan). Movies were recorded using EPU (Thermo Fisher Scientific,TFS) and AFIS (aberration-free image shift) in super-resolution mode at 64,000 nominal magnification, corresponding to a calibrated pixel size of 0.685 Å with a defocus range of −1 to −5 µm and a total dose of 30 e/Å ^2^(see data collection parameters in Figure. S1B).

### Cryo-ET data processing

Motion correction, frame alignment, tilt series alignment using gold fiducials, and tomogram reconstruction were carried out using IMOD (Kremer *et al*., 1996). The datasets were binned by a factor of 2. From the reconstructed tomograms, ϕKp24 phages with clearly visible baseplates, tail fibers were picked for segmentation using IMOD. Visualization of data was performed by IMOD and Fiji (Schindelin et al., 2012).

### SPA reconstruction of the capsid

The capsid of phage ϕKp24 was reconstructed using Relion 3.1.2 (Scheres, 2012). MotionCorr (Zheng et al., 2017a) was used to correct for beam-induced particle movement. The contrast transfer function (CTF) was estimated by CTFFIND 4.1.18 (Rohou and Grigorieff, 2015). Initially, 61 capsid particles were manually picked for 2D classification to generate reference templates for auto-picking. The Relion auto-picking was then used to automatically pick 290,280 particles, which were binned by 4 for 2D classification during extraction. After multiple rounds of 2D classification, particles with false-positive and contaminating features were discarded resulting in a 28,090 particle dataset, that was then classified into 2 classes: empty capsids and full capsids. To balance the final resolution and speed of computation, the dataset was down-sampled 1.5x with a final pixel size of 2.055 Å, a particle box of 800 pixels, and a circular mask diameter of 1600 Å. The extracted particles were then subjected to a 3D initial model and classification. The class with the largest number of particles was used for final 3D refinement (This separates and removes partially DNA-filled capsids). Finally, we chose both empty-capsid class and full-capsid class to 3D refinement separately using I1 icosahedral symmetry, Final reconstructions were sharpened and locally filtered by Relion post-processing (workflow of 3D reconstruction, Figure. S3A).

The map resolution was estimated at the 0.143 criterion of the phase-randomization-corrected FSC curve calculated between two independently refined half-maps multiplied by a soft-edged solvent mask. The maps were displayed using UCSF ChimeraX (Goddard *et al*., 2018).

### SPA reconstruction of the tail

The ϕKp24 tail tube particle was defined as a 400-pixels segment of the tail tube connected to a full capsid. A subset of 346 particles was manually picked and then extracted and 2D classified. The highest proportion of the classes with clear boundaries was used as templates for final auto picking. The parameters for auto picking were optimized on a subset of 741 micrographs. After auto picking from the whole dataset, 3,114,071 particles were extracted and binned by 4. Multiple rounds of 2D classification were performed to remove particles with false positive and contaminating features. An initial model with imposed C1 symmetry was generated and three-dimensional refinement was performed using followed by a 3D classification where the particles were classified into four classes. Particles in the class with the largest number were selected re-extracted using unbinned data and 3D refined with C1 symmetry. Final reconstructions were sharpened and locally filtered in Relion post-processing (3D reconstruction workflow Figure. S8A). The local resolution was generated in ResMap (Kucukelbir et al., 2014).

### Molecular modeling

The 3D structures of ϕKp24 proteins were inferred using Alphafold v2.0 via the ColabFold (Jumper et al., 2021) notebook at: https://colab.research.google.com/github/sokrypton/ColabFold/blob/main/AlphaFold2.ipynb.

#### Capsid pentamer and hexamer modeling

Initial models of the capsid hexamer and pentamer assemblies were constructed by rigidly docking the AlphaFold2-predicted gp372 model into corresponding regions of the capsid cryo-EM map using ChimeraX. The assembled models were then refined separately using the cascade Molecular Dynamics Flexible Fitting (MDFF) protocol (Singharoy et al., 2016), which conducts sequential MDFF simulations to a series of gaussian-smoothed maps of increasing resolution, ending with the experimental map. A total of 5 x 1ns MDFF simulations were carried out for each assembly with symmetry restraints applied to the backbone atoms of each monomer. The resulting hexamer model was then docked into the densities surrounding the hexamer and pentamer assemblies to produce extended capsid models, which were each then subjected to an additional 5-ns MDFF simulation. All simulations were conducted using NAMD v2.14 (Phillips et al., 2005) and the CHARMM36 force field (Huang and Roux, 2013). MDFF simulations were performed in the NVT ensemble at 300 K and using generalized Born implicit solvent. All backbone atoms were fit to the density map using a coupling constant of 0.3 with additional harmonic restraints applied to prevent loss of secondary structure, chirality errors, and the formation of cis-peptide bonds. Additional parameters were taken as the defaults provided by the MDFF plugin in VMD (Humphrey et al., 1996).

#### Helical Tail and Sheath

The AlphaFold2-predicted gp118 structure was fitted rigidly into tail density map using ChimeraX, the sheath region was then flexibly fitted by interactive molecular dynamics simulation in ISOLDE (Croll, 2018). The model of inner tube was constructed using PDB ID 5W5F as a template (Zheng *et al*., 2017b). The model was then optimized in ISOLDE.

### Tail fiber analysis using machine learning

A 100-layer mixed-scale dense neural network (Pelt and Sethian, 2018) was trained to detect phage ϕKp24 tail fibers in reconstructed tomograms. For training, manual segmentations of the tail fibers of seven phages were used, in addition to five manually selected regions without any fibers present. The ADAM algorithm (Kingma and Ba, 2014) was used to minimize the cross entropy loss, using random rotations and flips for data augmentation. Training was stopped after no significant improvement in the loss was observed, resulting in a training time of a few hours. Afterwards, the trained network was applied to all tomograms, resulting in 3D maps of tail fiber structures. To further analyze these maps, additional manual annotation was performed on the tomograms, indicating for each phage: (1) the center of the fiber structure, (2) the position of the head, and (3) whether the head was full, empty, or missing. Using this information, a 3D map of the fiber structure of each phage was extracted from the 3D fiber maps, resulting in individual maps for 89 phages with full heads, 338 phages with empty heads, and 181 phages with missing heads. The maps within each group were aligned to each other using both the manually annotated position information and subsequent automatic alignment by maximizing cross-correlation. The aligned maps were then averaged to produce ‘canonical’ fiber shapes for each group. The extracted structures and averaged structures were displayed using Paraview (Ahrens et al., 2005).

## QUANTIFICATION AND STATISTICAL ANALYSIS

The statistical analysis for each experiment is described in the figure legends.

## REFERENCES

Ahrens, J., Geveci, B., and Law, C. (2005). Paraview: An end-user tool for large data visualization. The visualization handbook 717.

Aksyuk, A.A., Kurochkina, L.P., Fokine, A., Forouhar, F., Mesyanzhinov, V.V., Tong, L., and Rossmann, M.G. (2011). Structural conservation of the myoviridae phage tail sheath protein fold. Structure 19, 1885–1894. 10.1016/j.str.2011.09.012.

Aksyuk, A.A., Leiman, P.G., Kurochkina, L.P., Shneider, M.M., Kostyuchenko, V.A., Mesyanzhinov, V.V., and Rossmann, M.G. (2009). The tail sheath structure of bacteriophage T4: a molecular machine for infecting bacteria. The EMBO Journal 28, 821–829. 10.1038/emboj.2009.36.

Altschul, S.F., Gish, W., Miller, W., Myers, E.W., and Lipman, D.J. (1990). Basic local alignment search tool. Journal of molecular biology 215, 403–410.

Baek, M., DiMaio, F., Anishchenko, I., Dauparas, J., Ovchinnikov, S., Lee, G.R., Wang, J., Cong, Q., Kinch, L.N., and Schaeffer, R.D. (2021). Accurate prediction of protein structures and interactions using a three-track neural network. Science 373, 871–876.

Basler, M. (2015). Type VI secretion system: secretion by a contractile nanomachine. Philosophical Transactions of the Royal Society B: Biological Sciences 370, 20150021. 10.1098/rstb.2015.0021.

Bonilla, B.E., Costa, A.R., Rossum, T.V., Hagedoorn, S., Walinga, H., Xiao, M., Song, W., Haas, P.-J., Nobrega, F.L., and Brouns, S.J.J. (2021). Genomic characterization of four novel bacteriophages infecting the clinical pathogen Klebsiella pneumoniae. Cold Spring Harbor Laboratory.

Clemens, D.L., Ge, P., Lee, B.-Y., Horwitz, M.A., and Zhou, Z.H. (2015). Atomic structure of T6SS reveals interlaced array essential to function. Cell 160, 940–951.

Cornelissen, A., Ceyssens, P.-J., T’syen, J., Van Praet, H., Noben, J.-P., Shaburova, O.V., Krylov, V.N., Volckaert, G., and Lavigne, R. (2011). The T7-related Pseudomonas putida phage φ15 displays virion-associated biofilm degradation properties. PloS one 6, e18597.

Cramer, P. (2021). AlphaFold2 and the future of structural biology. Nature Structural & Molecular Biology 28, 704–705. 10.1038/s41594-021-00650-1.

Croll, T.I. (2018). ISOLDE: a physically realistic environment for model building into low-resolution electron-density maps. Acta Crystallographica Section D: Structural Biology 74, 519–530.

Duda, R.L., and Teschke, C.M. (2019). The amazing HK97 fold: versatile results of modest differences. Current opinion in virology 36, 9–16.

Eskenazi, A., Lood, C., Wubbolts, J., Hites, M., Balarjishvili, N., Leshkasheli, L., Askilashvili, L., Kvachadze, L., van Noort, V., Wagemans, J., et al. (2022). Combination of pre-adapted bacteriophage therapy and antibiotics for treatment of fracture-related infection due to pandrug-resistant Klebsiella pneumoniae. Nature Communications 13, 302. 10.1038/s41467-021-27656-z.

Finn, R.D., Clements, J., and Eddy, S.R. (2011). HMMER web server: interactive sequence similarity searching. Nucleic acids research 39, W29–W37.

Follador, R., Heinz, E., Wyres, K.L., Ellington, M.J., Kowarik, M., Holt, K.E., and Thomson, N.R. (2016). The diversity of Klebsiella pneumoniae surface polysaccharides. Microbial genomics 2.

Gabler, F., Nam, S.Z., Till, S., Mirdita, M., Steinegger, M., Söding, J., Lupas, A.N., and Alva, V. (2020). Protein sequence analysis using the MPI bioinformatics toolkit. Current Protocols in Bioinformatics 72, e108.

Gan, L., Speir, J.A., Conway, J.F., Lander, G., Cheng, N., Firek, B.A., Hendrix, R.W., Duda, R.L., Liljas, L., and Johnson, J.E. (2006). Capsid conformational sampling in HK97 maturation visualized by X-ray crystallography and cryo-EM. Structure 14, 1655–1665.

Ge, P., Scholl, D., Leiman, P.G., Yu, X., Miller, J.F., and Zhou, Z.H. (2015). Atomic structures of a bactericidal contractile nanotube in its pre- and postcontraction states. Nat Struct Mol Biol 22, 377–382. 10.1038/nsmb.2995.

Goddard, T.D., Huang, C.C., Meng, E.C., Pettersen, E.F., Couch, G.S., Morris, J.H., and Ferrin, T.E. (2018). UCSF ChimeraX: Meeting modern challenges in visualization and analysis. Protein Science 27, 14–25.

Hagen, W.J., Wan, W., and Briggs, J.A. (2017). Implementation of a cryo-electron tomography tilt-scheme optimized for high resolution subtomogram averaging. Journal of structural biology 197, 191–198.

Helgstrand, C., Wikoff, W.R., Duda, R.L., Hendrix, R.W., Johnson, J.E., and Liljas, L. (2003). The refined structure of a protein catenane: the HK97 bacteriophage capsid at 3.44 Å resolution. Journal of molecular biology 334, 885–899.

Hendrix, R.W. (2009). Jumbo Bacteriophages. In Lesser Known Large dsDNA Viruses, (Springer Berlin Heidelberg), pp. 229–240. 10.1007/978-3-540-68618-7_7.

Heymann, J.B., Bartho, J.D., Rybakova, D., Venugopal, H.P., Winkler, D.C., Sen, A., Hurst, M.R., and Mitra, A.K. (2013). Three-dimensional structure of the toxin-delivery particle antifeeding prophage of Serratia entomophila. Journal of Biological Chemistry 288, 25276–25284.

Huang, L., and Roux, B. (2013). Automated force field parameterization for nonpolarizable and polarizable atomic models based on ab initio target data. Journal of chemical theory and computation 9, 3543–3556.

Humphrey, W., Dalke, A., and Schulten, K. (1996). VMD: visual molecular dynamics. Journal of molecular graphics 14, 33–38.

Jumper, J., Evans, R., Pritzel, A., Green, T., Figurnov, M., Ronneberger, O., Tunyasuvunakool, K., Bates, R., Žídek, A., and Potapenko, A. (2021). Highly accurate protein structure prediction with AlphaFold. Nature 596, 583–589.

Kamiya, R., Uchiyama, J., Matsuzaki, S., Murata, K., Iwasaki, K., and Miyazaki, N. (2021). Acid-stable capsid structure of Helicobacter pylori bacteriophage KHP30 by single-particle cryoelectron microscopy. Structure.

Kelley, L.A., Mezulis, S., Yates, C.M., Wass, M.N., and Sternberg, M.J. (2015). The Phyre2 web portal for protein modeling, prediction and analysis. Nature protocols 10, 845–858.

King, J., and Mykolajewyoz, N. (1973). Bacteriophage T4 tail assembly: proteins of the sheath, core and baseplate. Journal of molecular biology 75, 339–358.

Kingma, D.P., and Ba, J. (2014). Adam: A method for stochastic optimization. arXiv preprint arXiv:1412.6980.

Ko, K.S. (2017). The contribution of capsule polysaccharide genes to virulence of Klebsiella pneumoniae. Virulence 8, 485–486.

Kremer, J.R., Mastronarde, D.N., and McIntosh, J.R. (1996). Computer visualization of three-dimensional image data using IMOD. Journal of structural biology 116, 71–76.

Kucukelbir, A., Sigworth, F.J., and Tagare, H.D. (2014). Quantifying the local resolution of cryo-EM density maps. Nature methods 11, 63–65.

Kudryashev, M., Wang, R.Y.-R., Brackmann, M., Scherer, S., Maier, T., Baker, D., DiMaio, F., Stahlberg, H., Egelman, E.H., and Basler, M. (2015). Structure of the type VI secretion system contractile sheath. Cell 160, 952–962.

Latka, A., Leiman, P.G., Drulis-Kawa, Z., and Briers, Y. (2019). Modeling the architecture of depolymerase-containing receptor binding proteins in Klebsiella phages. Frontiers in microbiology 10, 2649.

Latka, A., Maciejewska, B., Majkowska-Skrobek, G., Briers, Y., and Drulis-Kawa, Z. (2017). Bacteriophage-encoded virion-associated enzymes to overcome the carbohydrate barriers during the infection process. Applied microbiology and biotechnology 101, 3103–3119.

Leiman, P.G., Basler, M., Ramagopal, U.A., Bonanno, J.B., Sauder, J.M., Pukatzki, S., Burley, S.K., Almo, S.C., and Mekalanos, J.J. (2009). Type VI secretion apparatus and phage tail-associated protein complexes share a common evolutionary origin. Proceedings of the National Academy of Sciences 106, 4154–4159.

Leiman, P.G., and Shneider, M.M. (2012). Contractile tail machines of bacteriophages. Viral molecular machines, 93–114.

Maghsoodi, A., Chatterjee, A., Andricioaei, I., and Perkins, N.C. (2019). How the phage T4 injection machinery works including energetics, forces, and dynamic pathway. Proceedings of the National Academy of Sciences 116, 25097–25105. 10.1073/pnas.1909298116.

Mastronarde, D.N. (2005). Automated electron microscope tomography using robust prediction of specimen movements. Journal of structural biology 152, 36–51.

McGreevy, R., Teo, I., Singharoy, A., and Schulten, K. (2016). Advances in the molecular dynamics flexible fitting method for cryo-EM modeling. Methods 100, 50–60.

Murray, C.J., Ikuta, K.S., Sharara, F., Swetschinski, L., Robles Aguilar, G., Gray, A., Han, C., Bisignano, C., Rao, P., Wool, E., et al. (2022). Global burden of bacterial antimicrobial resistance in 2019: a systematic analysis. The Lancet. 10.1016/s0140-6736(21)02724-0.

Pan, Y.-J., Lin, T.-L., Chen, C.-C., Tsai, Y.-T., Cheng, Y.-H., Chen, Y.-Y., Hsieh, P.-F., Lin, Y.-T., and Wang, J.-T. (2017). Klebsiella phage ΦK64-1 encodes multiple depolymerases for multiple host capsular types. Journal of virology 91, e02457–02416.

Pelt, D.M., and Sethian, J.A. (2018). A mixed-scale dense convolutional neural network for image analysis. Proceedings of the National Academy of Sciences 115, 254–259.

Phillips, J.C., Braun, R., Wang, W., Gumbart, J., Tajkhorshid, E., Villa, E., Chipot, C., Skeel, R.D., Kale, L., and Schulten, K. (2005). Scalable molecular dynamics with NAMD. Journal of computational chemistry 26, 1781–1802.

Plattner, M., Shneider, M.M., Arbatsky, N.P., Shashkov, A.S., Chizhov, A.O., Nazarov, S., Prokhorov, N.S., Taylor, N.M., Buth, S.A., and Gambino, M. (2019). Structure and function of the branched receptor-binding complex of bacteriophage CBA120. Journal of molecular biology 431, 3718–3739.

Prokhorov, N.S., Riccio, C., Zdorovenko, E.L., Shneider, M.M., Browning, C., Knirel, Y.A., Leiman, P.G., and Letarov, A.V. (2017). Function of bacteriophage G7C esterase tailspike in host cell adsorption. Molecular microbiology 105, 385–398.

Records, A.R. (2011). The type VI secretion system: a multipurpose delivery system with a phage-like machinery. Molecular plant-microbe interactions 24, 751–757.

Rohou, A., and Grigorieff, N. (2015). CTFFIND4: Fast and accurate defocus estimation from electron micrographs. Journal of structural biology 192, 216–221.

Rosenthal, P.B., and Henderson, R. (2003). Optimal determination of particle orientation, absolute hand, and contrast loss in single-particle electron cryomicroscopy. Journal of molecular biology 333, 721–745.

Ryan, K.J., and Ray, C.G. (2004). Medical microbiology. McGraw Hill 4, 370.

Scheres, S.H. (2012). RELION: implementation of a Bayesian approach to cryo-EM structure determination. Journal of structural biology 180, 519–530.

Schindelin, J., Arganda-Carreras, I., Frise, E., Kaynig, V., Longair, M., Pietzsch, T., Preibisch, S., Rueden, C., Saalfeld, S., and Schmid, B. (2012). Fiji: an open-source platform for biological-image analysis. Nature methods 9, 676–682.

Schwarzer, D., Buettner, F.F., Browning, C., Nazarov, S., Rabsch, W., Bethe, A., Oberbeck, A., Bowman, V.D., Stummeyer, K., and Mühlenhoff, M. (2012). A multivalent adsorption apparatus explains the broad host range of phage phi92: a comprehensive genomic and structural analysis. Journal of virology 86, 10384–10398.

Seul, A., Müller, J.J., Andres, D., Stettner, E., Heinemann, U., and Seckler, R. (2014). Bacteriophage P22 tailspike: structure of the complete protein and function of the interdomain linker. Acta Crystallographica Section D: Biological Crystallography 70, 1336–1345.

Šimoliūnas, E., Kaliniene, L., Truncaitė , L., Zajanč kauskaitė , A., Staniulis, J., Kaupinis, A., Ger, M., Valius, M., and Meškys, R. (2013). Klebsiella phage vB_KleM-RaK2—A giant singleton virus of the family Myoviridae. PLoS One 8, e60717.

Singharoy, A., Teo, I., McGreevy, R., Stone, J.E., Zhao, J., and Schulten, K. (2016). Molecular dynamics-based refinement and validation for sub-5 Å cryo-electron microscopy maps. Elife 5, e16105.

Suhanovsky, M.M., and Teschke, C.M. (2015). Nature׳ s favorite building block: Deciphering folding and capsid assembly of proteins with the HK97-fold. Virology 479, 487–497.

Tacconelli, E., Carrara, E., Savoldi, A., Harbarth, S., Mendelson, M., Monnet, D.L., Pulcini, C., Kahlmeter, G., Kluytmans, J., and Carmeli, Y. (2018). Discovery, research, and development of new antibiotics: the WHO priority list of antibiotic-resistant bacteria and tuberculosis. The Lancet Infectious Diseases 18, 318–327.

Weigele, P.R., Scanlon, E., and King, J. (2003). Homotrimeric, β-stranded viral adhesins and tail proteins. Journal of bacteriology 185, 4022–4030.

Wikoff, W.R., Liljas, L., Duda, R.L., Tsuruta, H., Hendrix, R.W., and Johnson, J.E. (2000). Topologically linked protein rings in the bacteriophage HK97 capsid. Science 289, 2129–2133.

Yamada, T., Satoh, S., Ishikawa, H., Fujiwara, A., Kawasaki, T., Fujie, M., and Ogata, H. (2010). A jumbo phage infecting the phytopathogen Ralstonia solanacearum defines a new lineage of the Myoviridae family. Virology 398, 135–147.

Yan, J., Mao, J., and Xie, J. (2014). Bacteriophage polysaccharide depolymerases and biomedical applications. BioDrugs 28, 265–274.

Yuan, Y., and Gao, M. (2017). Jumbo bacteriophages: an overview. Frontiers in microbiology 8, 403.

Zheng, S.Q., Palovcak, E., Armache, J.-P., Verba, K.A., Cheng, Y., and Agard, D.A. (2017a). MotionCor2: anisotropic correction of beam-induced motion for improved cryo-electron microscopy. Nature methods 14, 331–332.

Zheng, W., Wang, F., Taylor, N.M., Guerrero-Ferreira, R.C., Leiman, P.G., and Egelman, E.H. (2017b). Refined cryo-EM structure of the T4 tail tube: exploring the lowest dose limit. Structure 25, 1436–1441. e1432.

Zimmermann, L., Stephens, A., Nam, S.-Z., Rau, D., Kübler, J., Lozajic, M., Gabler, F., Söding, J., Lupas, A.N., and Alva, V. (2018). A completely reimplemented MPI bioinformatics toolkit with a new HHpred server at its core. Journal of molecular biology 430, 2237–2243.

