## supplementary files for "High resolution reconstruction of a Jumbo bacteriophage infecting capsulated bacteria using hyperbranched tail fibers"

A

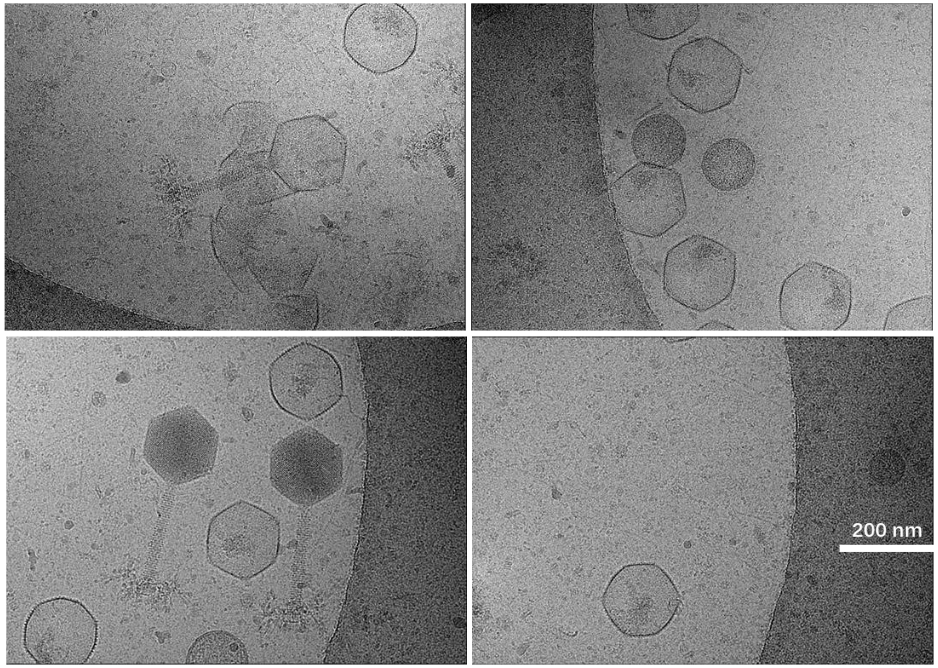


B

| | |
|---|---|
| Microscope | Krios 1 |
| Voltage (kV) | 300 |
| CS (mm) | 2.7 |
| Nominal magnification | SA 64000 |
| C1 | 2000 |
| Spot size | 6 |
| C2 aperture | 50 |
| Illumination area | 1.3 um |
| Objective aperture | 100 |

| | |
|---|---|
| Data collection software | EPU 2.8.1, AFIS |
| Detector | Gatan K3 |
| Energy filter (eV) | 20 |
| Detector mode | SuperRes Counted, non-CDS |
| Pixel size (A) | 0.685 |
| Total dose (sample e/A$^2$ ) | 30 |
| Exposure time (s) | 1.6 |
| Number of fractions | 30 |
| Data format | tiff lzw, non gain-normalized |
| Exposure per hole | 4 |
| Focus range (microns) | -1 to -5 |
| Repeated focus distance | 10 microns |

**FIG S1** (A) The Phage ϕKp24 representative micrographs. (B) Single-particle data collection parameters

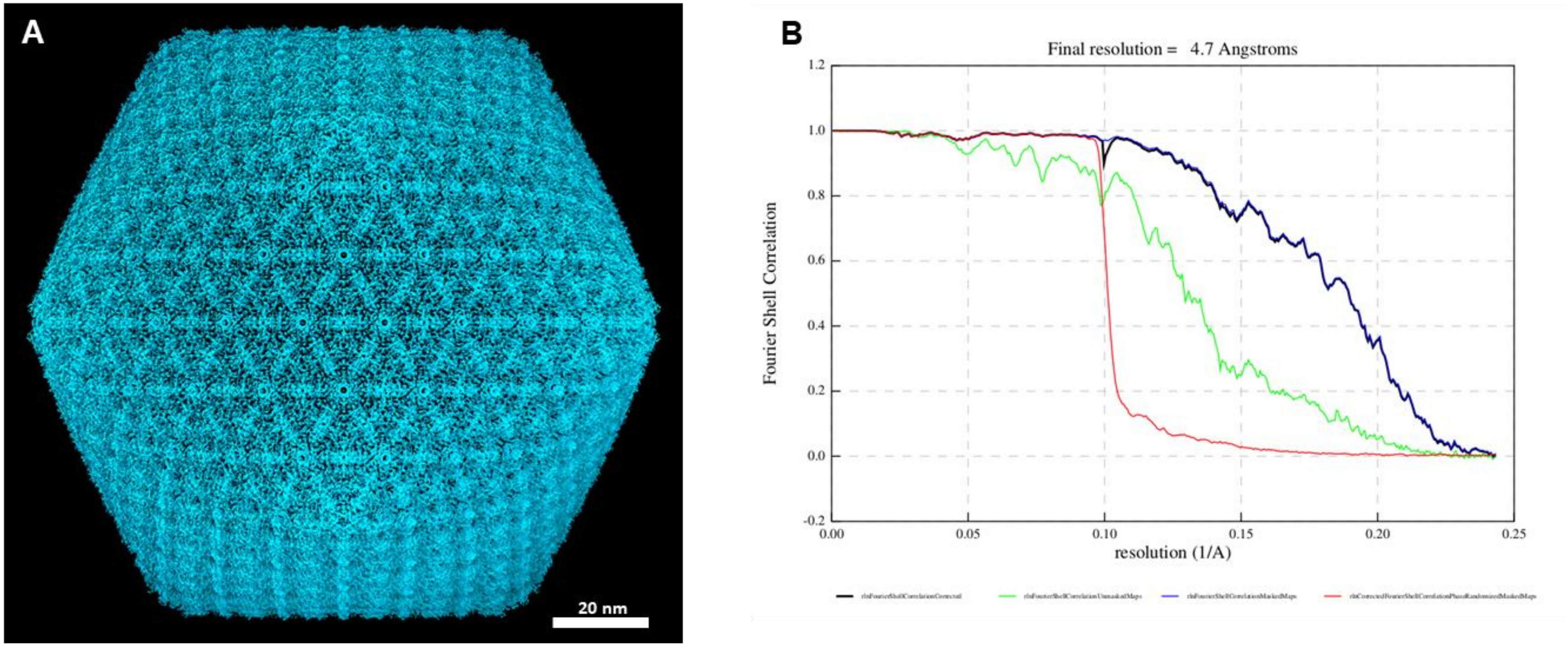


**FIG S2** (A) The Phage ϕKp24 full-capsid reconstruction (EMD-14356, cyan) showed in ChimeraX. (B) The "gold-standard" $FSC_{0.143}$ criterion shows the full capsid was reconstructed to 4.7Å resolution after polishing.

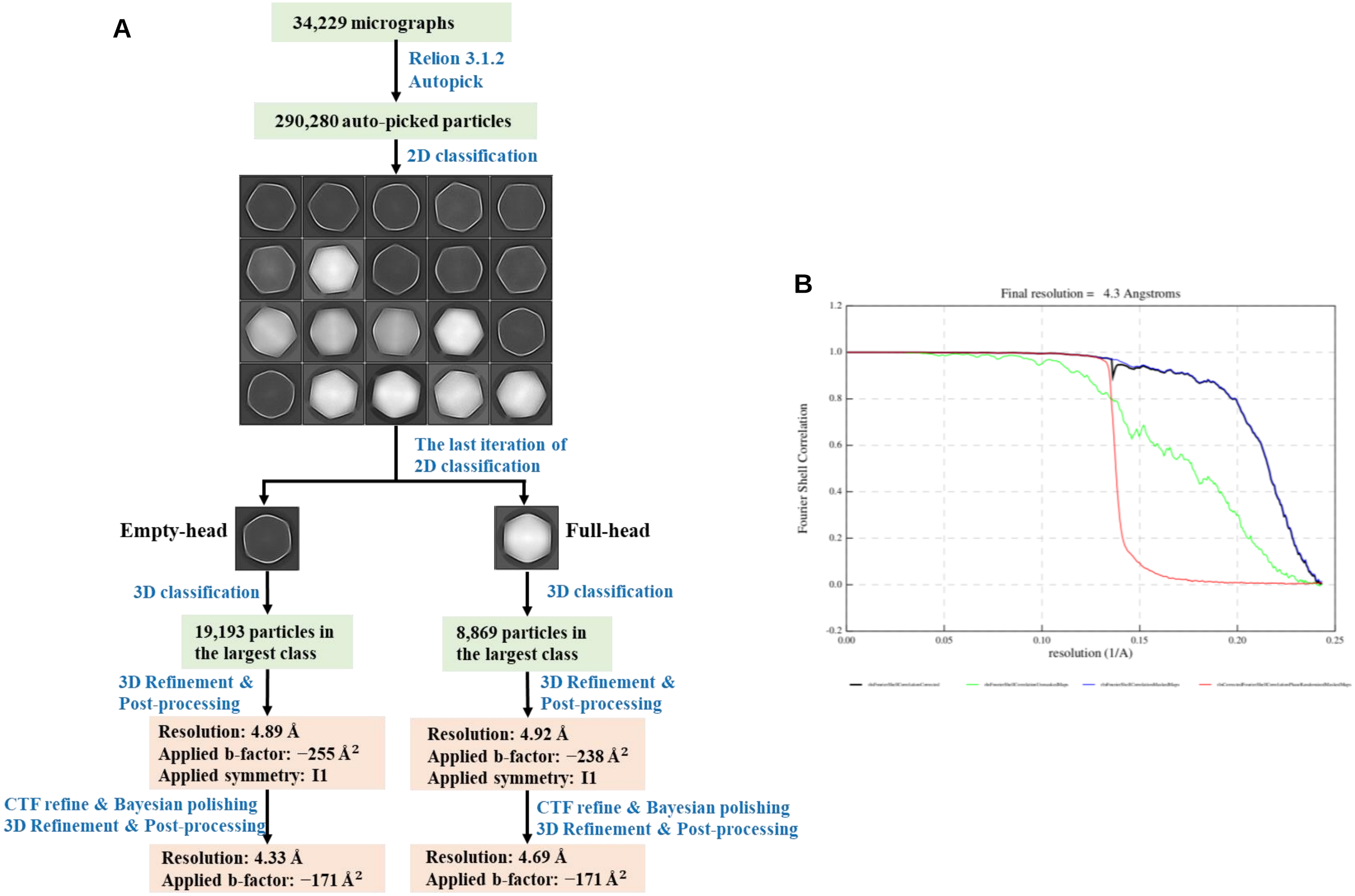


**FIG S3** (A) Workflow for the cryo-EM 3D reconstruction of both empty capsid and full capsid of Phage ϕKp24. The boxsize of representative 2D classes is 800 pixels, the mask diameter is 1600 Å. (B) The "gold-standard" $FSC_{0.143}$ criterion shows the empty capsid was reconstructed to 4.3Å resolution after polishing.

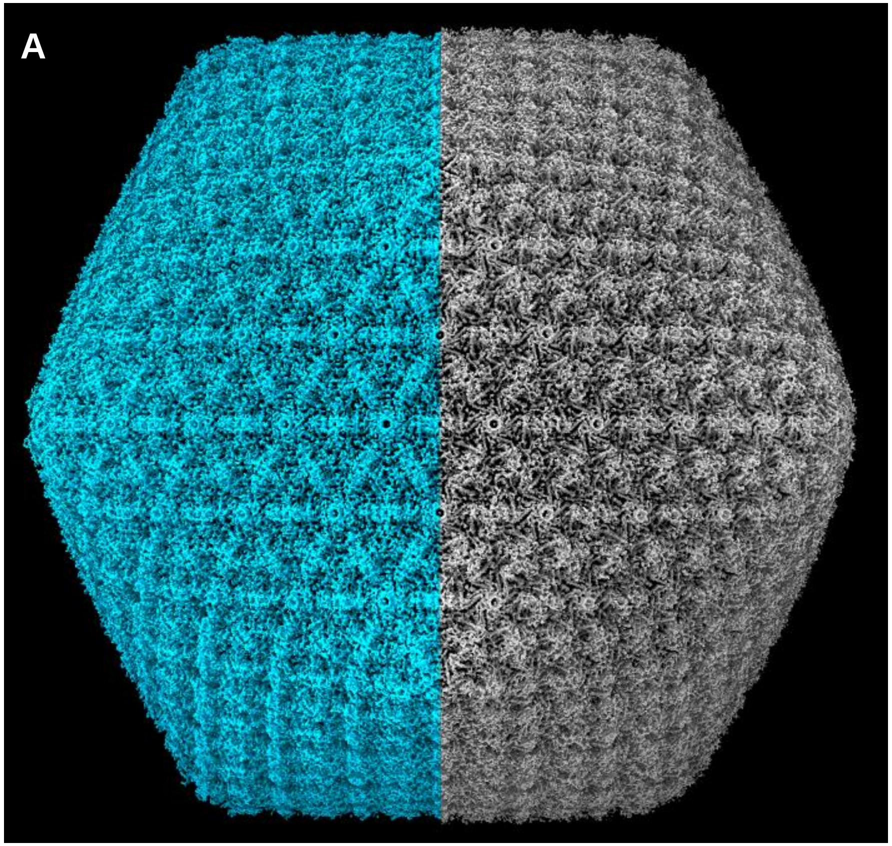


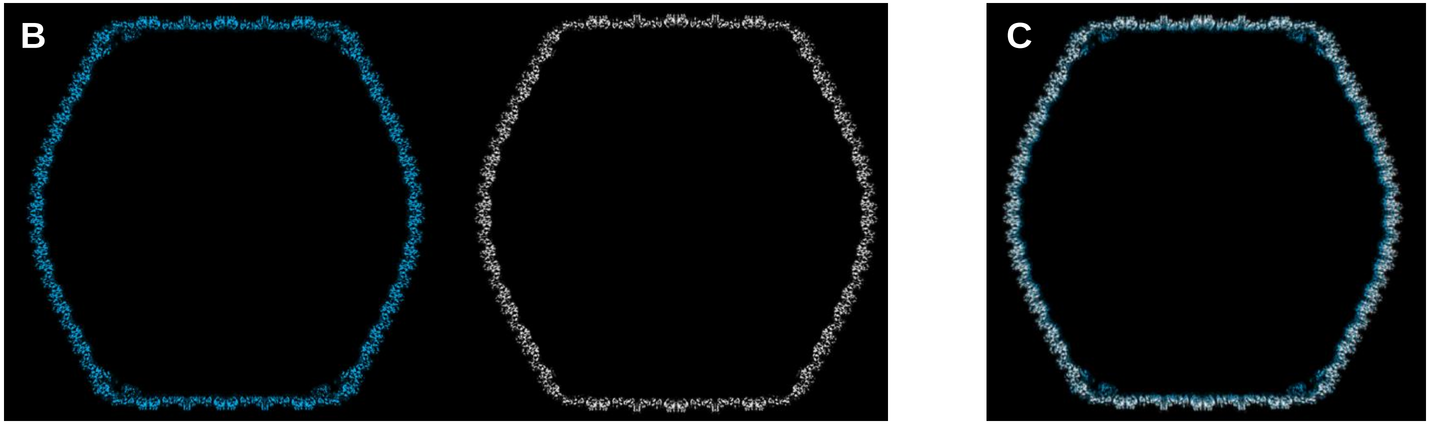


**FIG S4** (A) The comparation of phage ϕKp24 full capsid (cyan) and empty capsid (gray) showed in ChimeraX. The two capsids use the same organization. (B) Central cross-section of full capsid (cyan) and empty capsid (gray) viewed along an icosahedral two-fold axis. (C) The overlay view of two central cross-sections showed in B. The full capsid (cyan) is a little smaller than empty capsid (gray), which means the full-DNA capsid is more compact than the empty one.

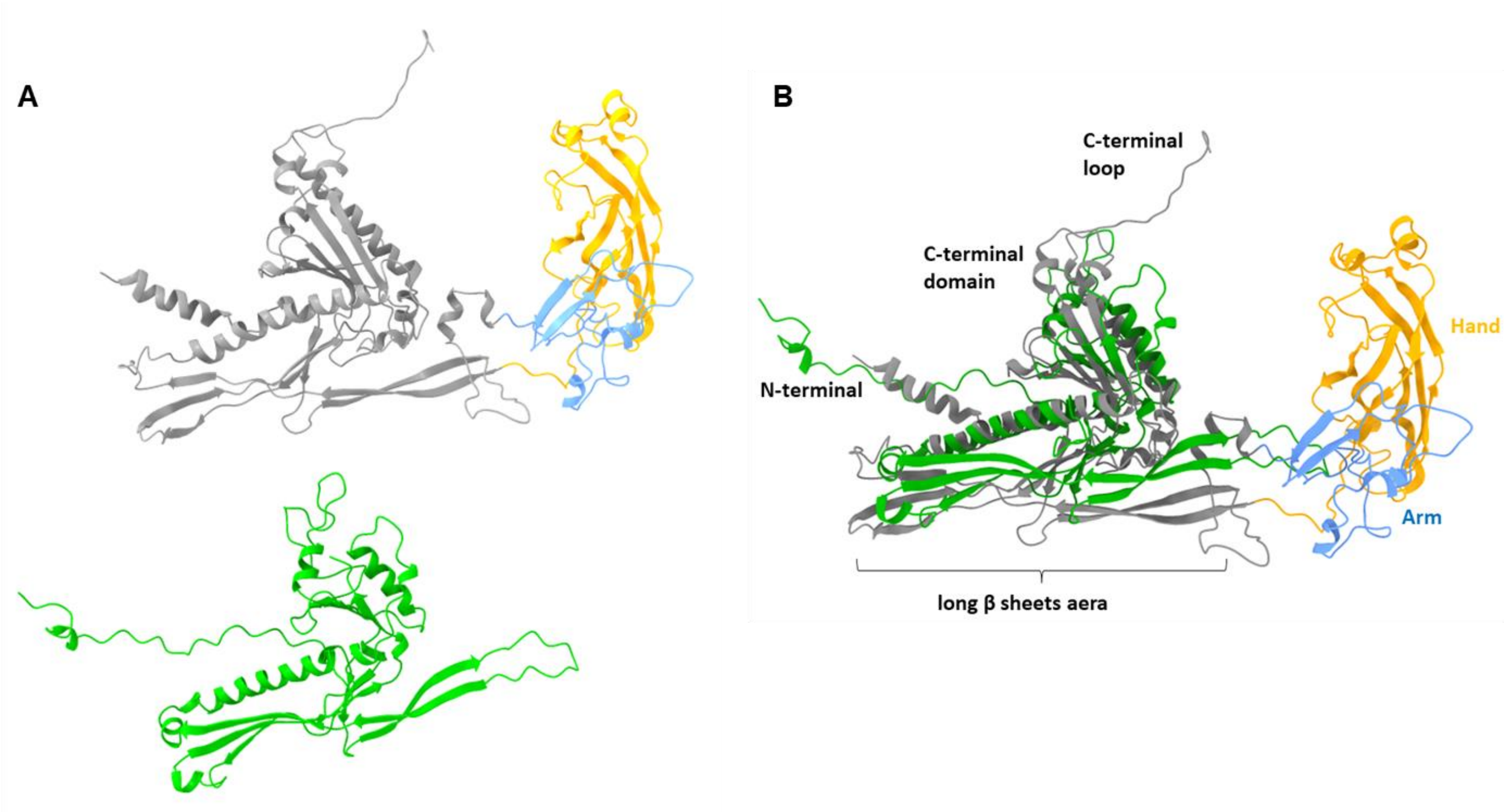


**FIG S5 .** (A) A separated view of the comparison between Phage ϕKp24's MCP gp372(gray, blue, orange) and the MCP (green) of HK97. (B) The triangle-part (gray) of Phage ϕKp24's MCP gp372 compared with MCP (green) of HK97. The triangle-part shows a similar core folds as same as phage HK97, except the long C-terminal loop, the N-terminal and C-terminal domain. The size of long β sheets aera is different.

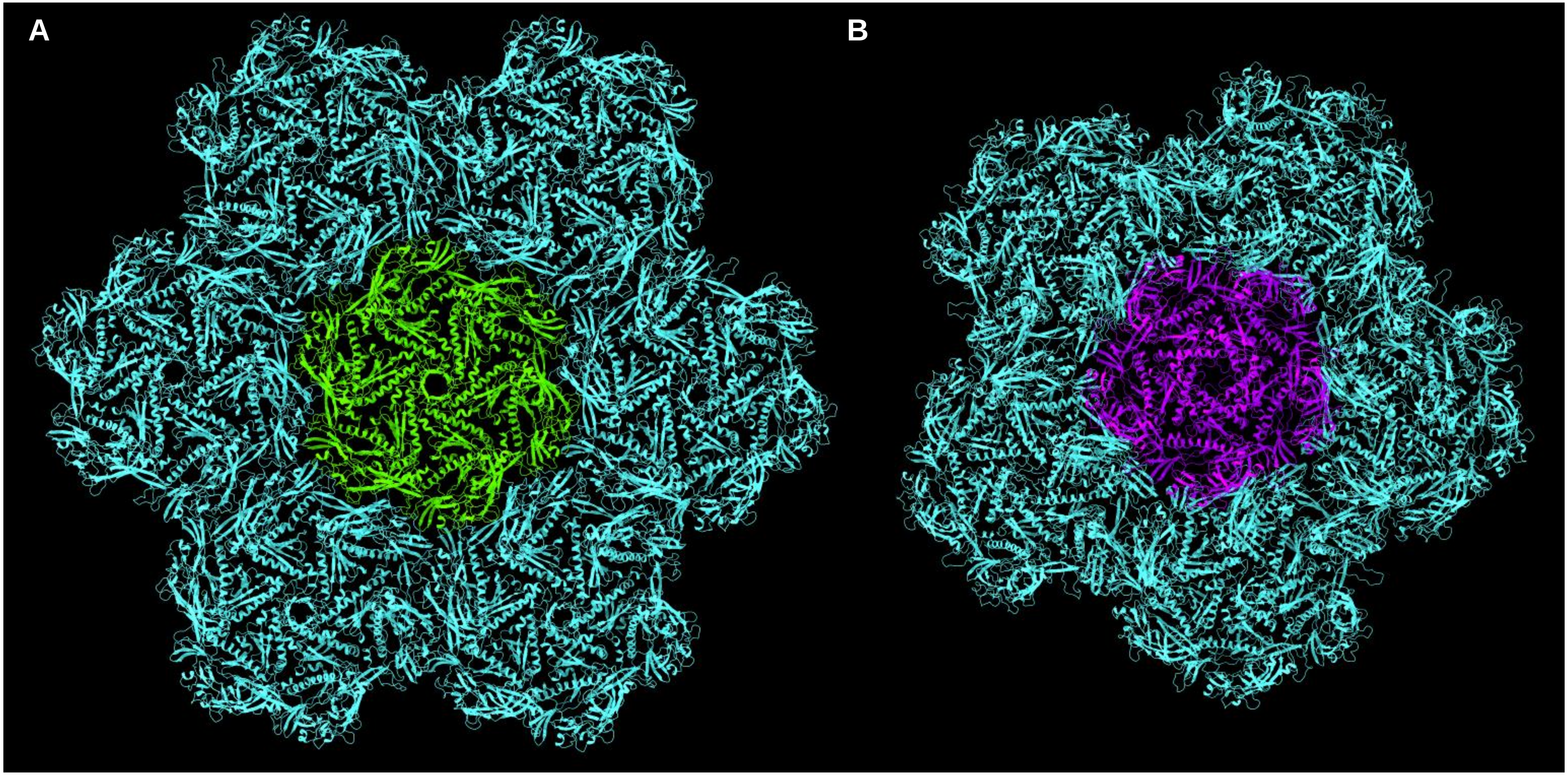


**FIG S6** (A) A hexamer surrounding with six hexamers. One hexamer consists of six gp372 proteins and the model was built by MDFF method. The center hexamer was colored in green, other hexamers were colored in cyan. (B) A pentamer surrounding with five hexamers. Pentamer consists of five gp372 proteins. The center pentamer was colored in purple and hexamers were colored in cyan.

**FIG S7** the lists of the putative inter-MCP contacts for the hexamer and pentamer.

| Hexamer contacts | | | | | |
|---|---|---|---|---|---|
| Triangle-triangle interactions, CW neighbor, same hexamer | | Triangle-triangle interactions, CCW neighbor, same hexamer | | HAND-HAND interactions, neighboring hexamer | |
| LYS,380 | GLU,220 | ASP,163 | ARG,558 | GLU,201 | LYS,210 |
| GLU,398 | ARG,190 | ARG,190 | GLU,398 | LYS,210 | GLU,201 |
| ARG,477 | GLU,442 | GLU,220 | LYS,380 | LYS,210 | GLU,251 |
| ARG,554 | GLU,313 | GLU,313 | ARG,554 | LYS,247 | ASP,260 |
| ARG,558 | ASP,163 | ASP,419 | ARG,583 | ASP,249 | ARG,264 |
| ARG,583 | ASP,419 | GLU,442 | ARG,477 | GLU,251 | LYS,210 |
| ARG,583 | ASP,587 | ASP,587 | ARG,583 | GLU,251 | ARG,264 |
| | | | | ASP,260 | LYS,247 |
| | | | | ASP,260 | ARG,264 |
| | | | | ARG,264 | ASP,249 |
| | | | | ARG,264 | GLU,251 |
| | | | | ARG,264 | ASP,260 |

| Pentamer contacts | | | | | |
|---|---|---|---|---|---|
| Triangle-triangle interactions, CW neighbor, same pentamer | | Triangle-triangle interactions, CCW neighbor, same pentamer | | HAND-HAND interactions, neighboring hexamer | |
| LYS,380 | GLU,220 | ASP,163 | ARG,558 | GLU,201 | LYS,210 |
| ARG,501 | ASP,448 | GLU,220 | LYS,380 | LYS,210 | GLU,201 |
| ARG,558 | ASP,163 | GLU,442 | ARG,583 | LYS,210 | GLU,251 |
| ARG,583 | GLU,442 | ASP,448 | ARG,501 | LYS,247 | ASP,260 |
| | | | | ASP,249 | ARG,264 |
| | | | | GLU,251 | LYS,210 |
| | | | | ASP,260 | LYS,247 |
| | | | | ARG,264 | ASP,249 |

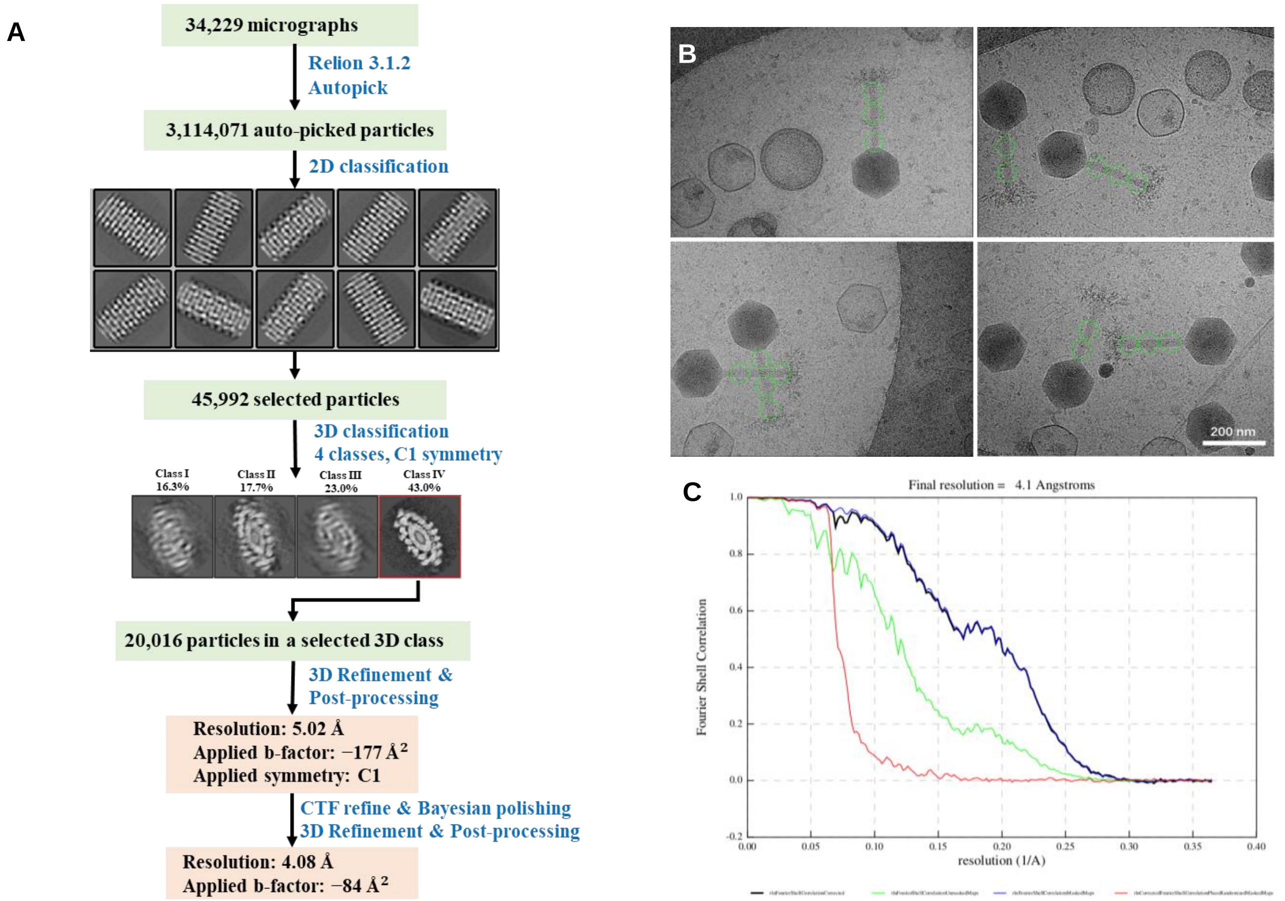


**FIG S8** (A) Workflow for the cryo-EM 3D reconstruction of helical tail structure. The box size of representative 2D classes is 400 pixels, the mask diameter is 540 Å. (B) A part of the helical tail connecting with full-capsid shows as a particle (green circle). (C) The "gold-standard" $FSC_{0.143}$ criterion shows the helical tail structure in its extended conformation has been determined to a resolution of 4.1 Å.

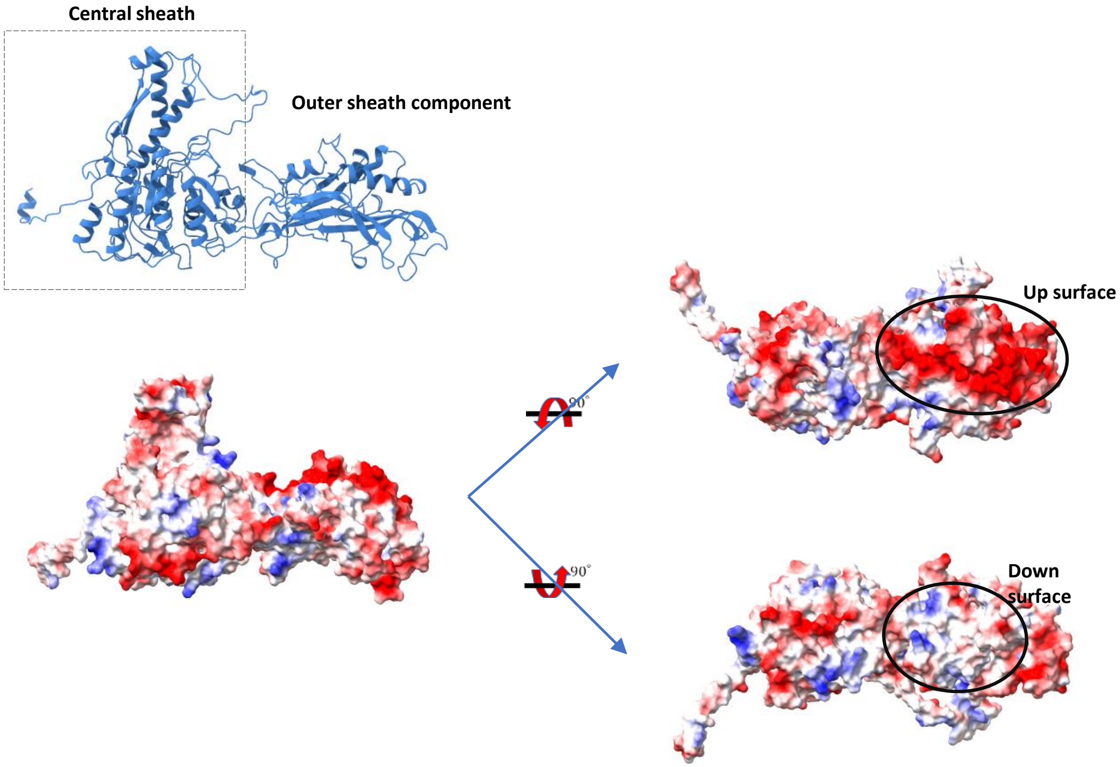


**FIG S9** The outer sheath component interactions caused by electrostatic forces.

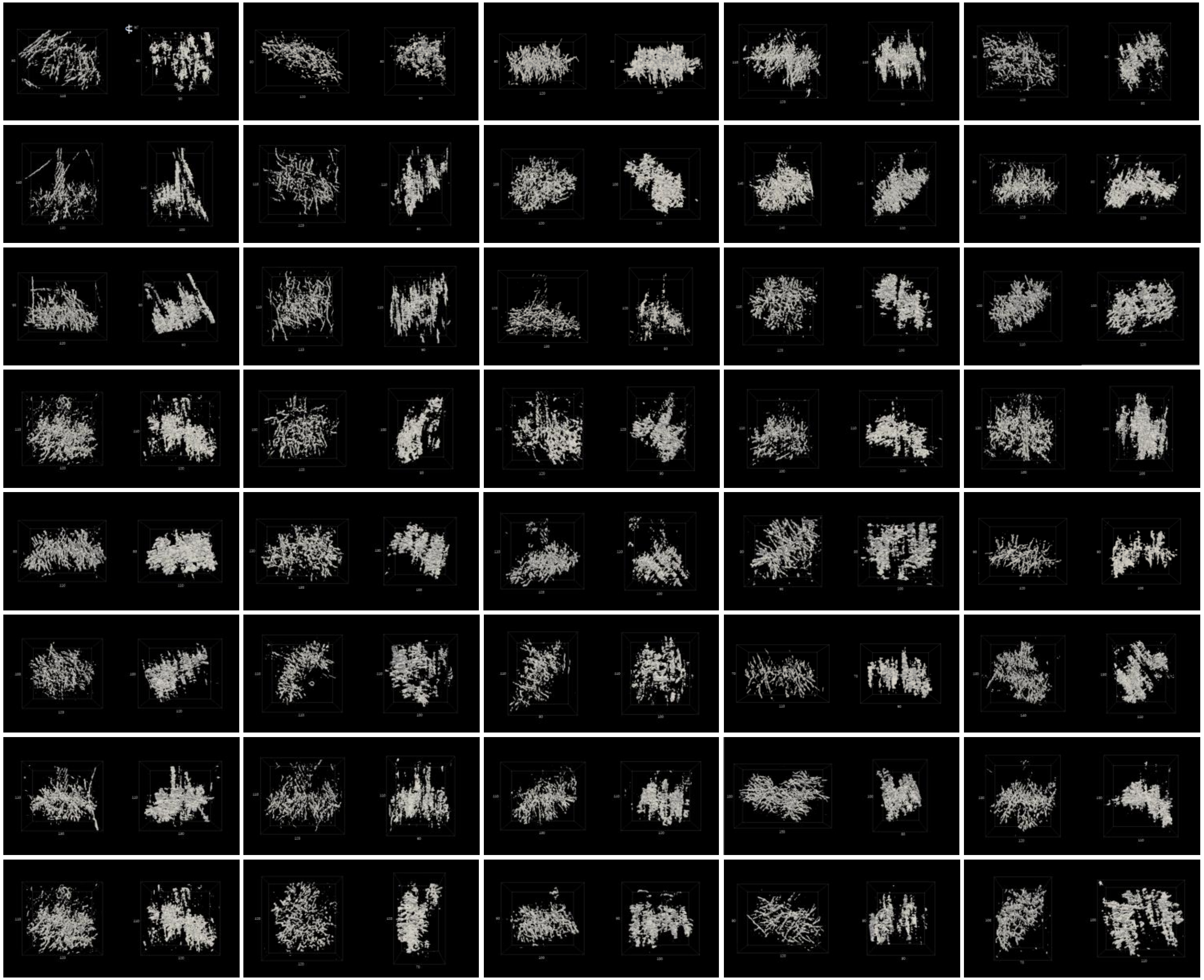

**FIG S10** 40 tail-fibers structures of pre-infection phages. Each right image shows the side view of the left structure after rotation by 90˚. Both the outer box and red line are scale bar with pixel unit. Pixel size is 1.312 nm.

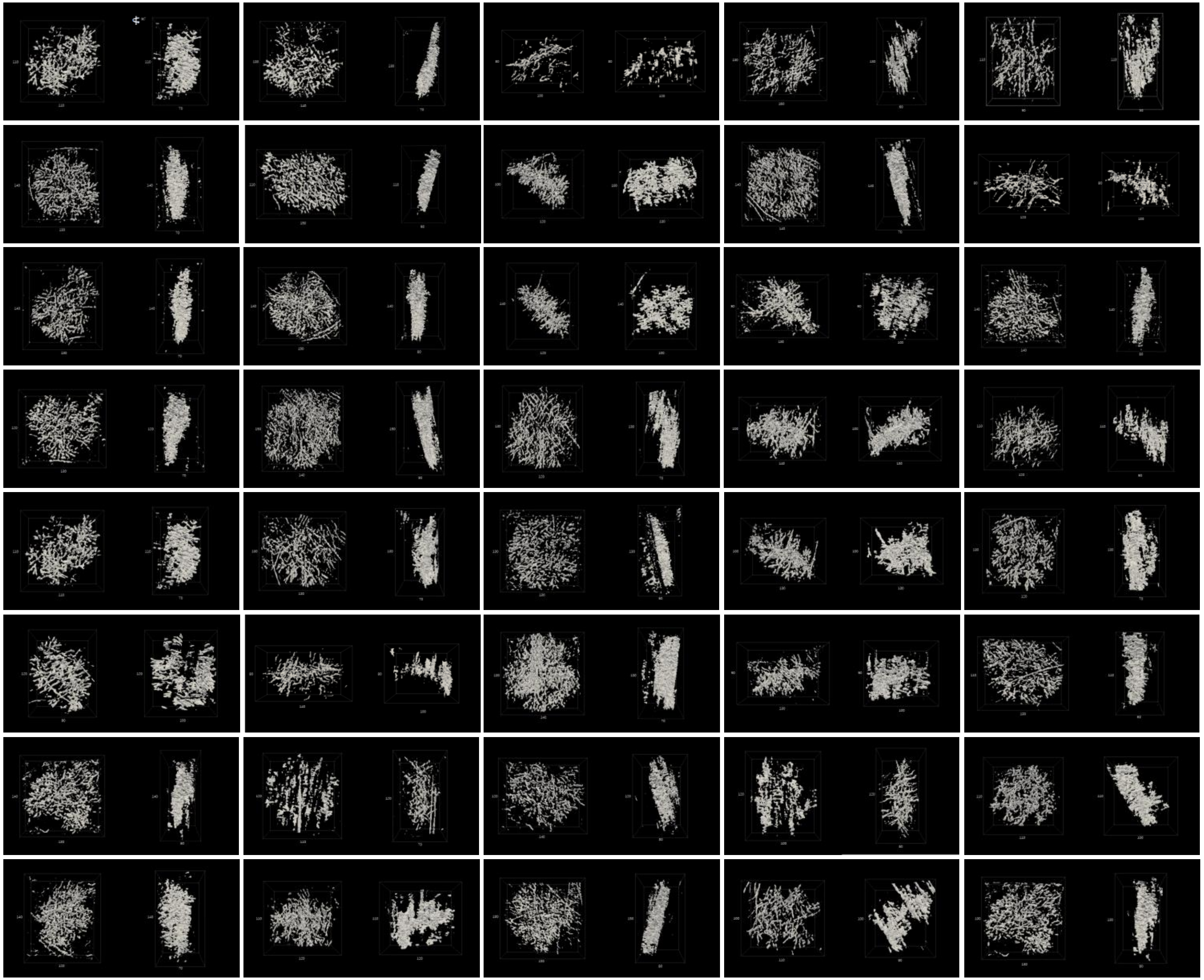

**FIG S11** 40 tail-fibers structures of post-infection phage attached to the cell. Each right image shows the side view of the left structure after rotation by 90˚. Both the outer box and red line are scale bar with pixel unit. Pixel size is 1.312 nm.

| Serotype | Strain | Phage ϕKp24 | | | | |
|---|---|---|---|---|---|---|
| | | Lytic activity | Capsule depolymerase-generated halo zone | E.O.P. | The titer of the phage at the terminal dilution | Predicted serotype specificity of depolymerases |
| K1 | CIP 52.144 | - | - | | | |
| K2 | CIP 52.145 | + | halo zone | 1 | 10^7 | gp196 (similar to gp43 KLPN1) |
| K3 | CIP 52.146 | - | - | | | |
| K4 | CIP 52.211 T | - | - | | | |
| K5 | CIP 52.212 | - | - | | | |
| K6 | CIP 52.213 | - | - | | | |
| K7 | CIP 52.205 | - | - | | | |
| K8 | CIP 52.206 | - | - | | | |
| K9 | CIP 52.207 | - | - | | | |
| K10 | CIP 52.214 | - | - | | | |
| K11 | CIP 52.215 | - | - | | | |
| K12 | CIP 52.216 | - | - | | | |
| K13 | CIP 52.217 | + | halo zone | 0.001 | 10^5 | |
| K14 | CIP 52.218 | - | - | | | |
| K15 | CIP 52.21 | - | - | | | |
| K16 | CIP 52.220 | - | - | | | |
| K17 | CIP 52.221 | - | - | | | |
| K18 | CIP 52.222 | - | - | | | |
| K19 | CIP 52.223 | + | halo zone | 0.01 | 10^5 | |
| K20 | CIP 52.224 | - | - | | | |
| K21 | CIP 52.225 | - | - | | | |
| K21 | CIP 52.969 | - | - | | | |
| K21 | CIP 52.358 | - | - | | | |
| K22 | CIP 52.226 | - | - | | | |
| K23 | CIP 52.228 | - | - | | | |
| K24 | CIP 52.229 | - | - | | | |
| K25 | CIP 52.230 | + | - | 0.01 | 10^5 | gp300 (similar to gp59 K64-1) |
| K26 | CIP 53.6 | - | - | | | |
| K27 | CIP 52.232 | - | - | | | |
| K28 | CIP 52.233 | - | - | | | |
| K29 | CIP 52.234 | - | - | | | |
| K30 | CIP 52.235 | - | - | | | |
| K31 | CIP 52.231 | - | - | | | |
| K32 | CIP 53.7 | - | - | | | |
| K33 | CIP 53.8 | - | - | | | |
| K34 | CIP 53.9 | - | - | | | |
| K35 | CIP 53.10 | + | halo zone | 1 | 10^7 | gp301 (similar to gp60 K64-1) |
| K36 | CIP 53.11 | - | - | | | |
| K37 | CIP 53.12 | - | - | | | |
| K38 | CIP 53.13 | - | - | | | |
| K39 | CIP 53.14 | - | - | | | |
| K40 | CIP 53.15 | - | - | | | |
| K41 | CIP 53.16 | - | - | | | |
| K42 | CIP 53.17 | - | - | | | |
| K43 | CIP 53.19 | - | - | | | |
| K44 | CIP 53.20 | - | - | | | |
| K45 | CIP 53.21 | - | - | | | |
| K46 | CIP 53.22 | + | halo zone | 1 | 10^7 | |
| K47 | CIP 53.23 | - | - | | | |
| K48 | CIP 53.24 | - | - | | | |
| K49 | CIP 52.199 | - | - | | | |
| K50 | CIP 52.200 | - | - | | | |
| K51 | CIP 52.201 | - | - | | | |
| K52 | CIP 53.25 | - | - | | | |
| K53 | CIP 53.26 | - | - | | | |
| K54 | CIP 53.27 | - | - | | | |
| K55 | NCTC 9175 | - | - | | | |
| K56 | NCTC 9176 | - | - | | | |
| K57 | NCTC 9177 | - | - | | | |
| K58 | NCTC 9178 | - | - | | | |
| K59 | NCTC 9179 | - | - | | | |
| K60 | NCTC 9180 | - | - | | | |
| K61 | NCTC 9181 | + | halo zone | 0.1 | 10^6 | |
| K62 | NCTC 9182 | - | - | | | |
| K63 | NCTC 9183.1 | - | - | | | |
| | NCTC 9183.2 | - | - | | | |
| K64 | K6592 (HOST) | + | halo zone | 1 | 10^7 | |
| | NCTC 9184 | + | halo zone | 1 | 10^7 | |
| K64 | CIP 80.47 | - | | | | |
| K65 | NCTC 9185 | - | | | | |
| K66 | NCTC 9186 | - | | | | |
| K67 | NCTC 9187 | - | | | | |
| K68 | NCTC 9188 | - | | | | |
| K69 | NCTC 9189 | - | | | | |
| K70 | NCTC 10261 | - | | | | |
| K71 | NCTC 10262 | - | | | | |
| K72 | NCTC 10263 | - | | | | |
| K74 | NCTC 11355 | - | | | | |
| K79 | NCTC 11356 | - | | | | |
| K80 | NCTC 11357 | - | | | | |
| K81 | NCTC 11358 | + | | 1 | 10^-5 | |
| K82 | NCTC 11359 | - | | | | |

**FIG S12** Using the Klebsiella strains from K serotype collection to identify the specific capsule degrading enzymes produced by phage ϕKp24

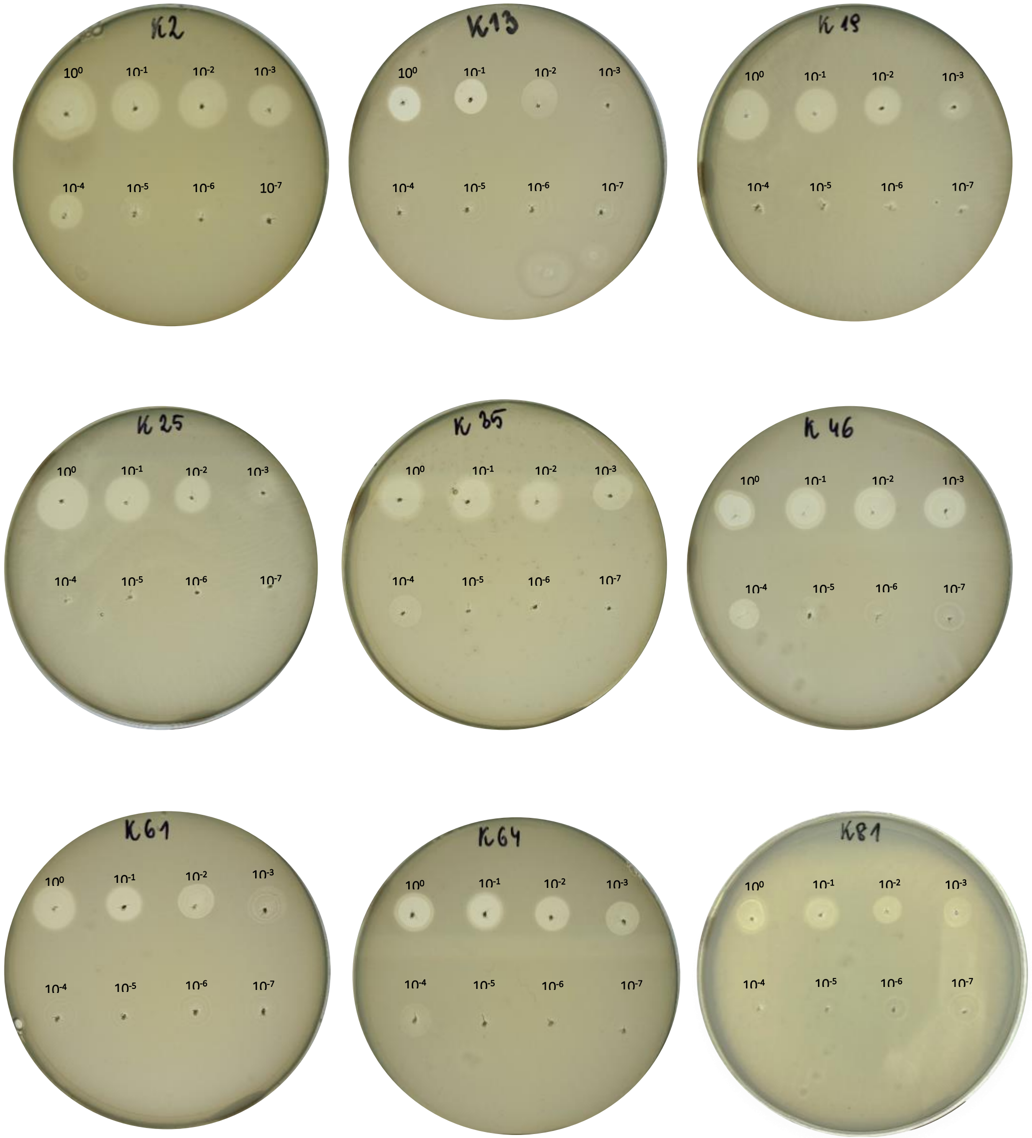


**FIG S13** A spot test of ϕKp24 phage on capsular serotypes of *Klebsiella pneumoniae*. 10 µl of each of a serial dilution ($10^0$-$10^{-7}$) of the ϕKp24 phage suspension ($10^7$ pfu/ml) was spotted onto bacterial lawn. The presence of plaques and capsule depolymerase-generated halo zone were observed after overnight incubation. The bacteriophage ϕKp24 can infect 9 capsular serotypes of *K. pneumoniae* **-** K2 (CIP 52.145), K13 (CIP 52.217), K19 (CIP 52.223), K25 (CIP 52.230), K35 (CIP 53.10), K46 (CIP 53.22), K61 (NCTC 9181), K64 (NCTC 9184) and K81 (NCTC 11358).

### Supplementary table 1

| NCBI | | | | RoseTTAfold | | | Phyre 2 | | | | | HHPred | | | | |
|---|---|---|---|---|---|---|---|---|---|---|---|---|---|---|---|---|
| Locus tag | Accession number | Definition | Length [aa] | Length to the α-helix [aa] | α-helix location [aa] | Confidence | Template | Alignment region | Confidence | %i.d. | β-helix | Alignment to | Probability [%] | Alignment region [aa] protein | Kp24 protein vs T4/CBA | Remark |
| vBKpMFBKp24_168 | QQV91997.1 | hypothetical protein | 881 | 194 | 195-211 | 0.66 | hydrolase, particle-associated glycoside hydrolase | 115-496 | 92.2 | 25 | YES | | | | | Group 3 |
| | | | | | | | hydrolase, alpha-1,3-glucanase | 187-646 | 96.1 | 19 | YES | | | | | |
| | | | | | | | hydrolase, exo-poly-alpha-d-galacturonosidase; | 177-646 | 97.2 | 18 | YES | | | | | |
| | | | | | | | hydrolase, pectate lyase | 176-649 | 97.6 | 14 | YES | | | | | |
| | | | | | | | single-stranded right-handed beta-helix, Pectin lyase-like, galacturonase | 189-574 | 96.3 | 17 | YES | | | | | |
| vBKpMFBKp24_196 | QQV92020.1 | putative tail fiber protein | 660 | 184 | 185-202 | 0.79 | viral protein, hydrolase, tailspike protein | 96-646 | 99.9 | 22 | YES | | | | | Group 3 |
| | | | | | | | hydrolase, glucan 1,3-beta-glucosidase | 173-656 | 100 | 19 | YES | | | | | |
| | | | | | | | hydrolase, exopolygalacturonase | 31-466 | 99.8 | 17 | YES | | | | | |
| vBKpMFBKp24_294 | QQV92014.1 | bifunctional tail protein | 689 | 311 | 312-322 | 0.60 | hydrolase | 290-518 | 98.3 | 19 | YES | | | | | Group 2 |
| | | | | | | | hydrolase, xylosidase | 293-532 | 98.2 | 16 | YES | | | | | |
| | | | | | | | hydrolase, exopolygalacturonase | 449-574 | 39.7 | 18 | YES | | | | | |
| vBKpMFBKp24_295 | QQV92015.1 | bifunctional tail protein | 679 | 316 | 317-325 | 0.60 | hydrolase, xylosidase | 295-612 | 98.6 | 16 | YES | | | | | Group 2 |
| vBKpMFBKp24_300 | QQV92023.1 | putative tail fiber protein | 598 | 142 | 143-155 | 0.64 | isomerase, poly(beta-d-mannuronate) c5 epimerase; | 234-373 | 80.8 | 19 | YES | | | | | Group 3 |
| | | | | | | | isomerase, poly(beta-d-mannuronate) c5 epimerase 4 | 146-373 | 37.3 | 21 | YES | | | | | |
| | | | | | | | single-stranded right-handed beta-helix, pectin lyase-like | 146-346 | 33.9 | 19 | YES | | | | | |
| vBKpMFBKp24_301 | QQV91995.1 | hypothetical protein | 915 | 289 | 290-305 | 0.48 | cell invasion, putative pectin lyase | 289-491 | 97.5 | 30 | YES | T4gp10 | 47.23 | 108-188 vs 150-246 | | Group 2 |
| | | | | | | | transferase, beta-1,3-glucanase | 289-485 | 92.5 | 24 | YES | T4gp10 | 44.39 | 128-188 vs 170-194 | | Presence T4gp10-like domain detected |
| | | | | | | | hydrolase, glucan 1,3-beta-glucosidase | 289-486 | 92.4 | 23 | YES | T4gp10 | 41.77 | 152-186 vs 347-384 | | 61-194 |
| | | | | | | | | | | | | CBA120gp163 | 98.87 | 111-189 vs 11-82 | | |
| | | | | | | | | | | | | CBA120gp165 | 99.77 | 61-194 vs 204-338 | | |
| | | | | | | | | | | | | CBA120gp165 | 94.05 | 115-189 vs 183-250 | | |
| vBKpMFBKp24_303 | QQV92019.1 | putative tail fiber protein | 661 | 99 | 100-120 | 0.71 | viral protein, hydrolase, tailspike protein | 21-301 | 99.4 | 25 | YES | | | | | Group 3 |
| | | | | | | | cell invasion, putative pectin lyase | 45-375 | 97.5 | 17 | YES | | | | | |
| | | | | | | | hydrolase, exopolygalacturonase | 33-355 | 97.7 | 14 | YES | | | | | |
| | | | | | | | single-stranded right-handed beta-helix, pectin lyase-like | 88-435 | 96.1 | 13 | YES | | | | | |
| | | | | | | | hydrolase, pectate lyase | 88-414 | 95.4 | 13 | YES | | | | | |
| vBKpMFBKp24_304 | QQV92004.1 | putative tail fiber protein | 755 | 178 | 179-199 | 0.64 | viral protein, hydrolase, tailspike protein | 91-424 | 100 | 22 | YES | | | | | Group 3 |
| | | | | | | | hydrolase, alpha-1,3-glucanase | 111-454 | 95.9 | 17 | YES | | | | | |
| | | | | | | | single-stranded right-handed beta-helix, pectin lyase-like, galacturonase | 162-480 | 96.3 | 17 | YES | | | | | |
| vBKpMFBKp24_306 | QQV91996.1 | putative tail fiber protein | 883 | 398 | 399-415 | 0.54 | hydrolase | 322-567 | 100 | 21 | YES | T4gp10 | 96.97 | 128-188 vs 283-248 | | Group 1 |
| | | | | | | | hydrolase, glucan 1,3-beta-glucosidase | 388-557 | 61.1 | 18 | YES | T4gp10 | 75.19 | 215-274 vs 169-247 | | Presence T4gp10-like domain detected |
| | | | | | | | single-stranded right-handed beta-helix, pectin lyase-like | 385-560 | 58.2 | 15 | YES | T4gp10 | 55.8 | 141-188 vs 305-387 | | 64-275 |
| | | | | | | | lyase | 391-557 | 53 | 18 | YES | CBA120gp163 | 99.65 | 111-271 vs 10-155 | | |
| | | | | | | | | | | | | CBA120gp163 | 98.3 | 64-185 vs 38-156 | | |
| | | | | | | | | | | | | CBA120gp163 | 86.5 | 207-274 vs 19-81 | | |
| | | | | | | | | | | | | CBA120gp165 | 99.89 | 118-286 vs 185-343 | | |
| | | | | | | | | | | | | CBA120gp165 | 99.59 | 61-188 vs 204-333 | | |
| | | | | | | | | | | | | CBA120gp165 | 95.83 | 203-275 vs 183-250 | | |
| vBKpMFBKp24_307 | QQV92007.1 | hypothetical protein | 742 | 157 | 158-170 | 0.69 | viral protein, preneck appendage protein | 131-692 | 100 | 18 | YES | | | | | Group 3 |
| | | | | | | | lyase, K5 lyase | 147-531 | 97.8 | 22 | YES | | | | | |
| | | | | | | | hydrolase, glycoside hydrolase | 144-508 | 100 | 20 | YES | | | | | |
| | | | | | | | hydrolase, alpha-1,3-glucanase | 62-539 | 100 | 14 | YES | | | | | |
| vBKpMFBKp24_308 | QQV92009.1 | hypothetical protein | 737 | 174 | 175-191 | 0.55 | hydrolase, endopolygalacturonase | 172-520 | 91.7 | 18 | YES | | | | | Group 3 |
| | | | | | | | single-stranded right-handed beta-helix, pectin lyase-like, galacturonase | 175-508 | 90.3 | 15 | YES | | | | | |
| | | | | | | | cell invasion, putative pectin lyase | 181-368 | 94.7 | 22 | YES | | | | | |
| | | | | | | | hydrolase, endo-xylogalacturonan hydrolase | 175-450 | 84.5 | 20 | YES | | | | | |
| vBKpMFBKp24_309 | QQV92005.1 | hypothetical protein | 751 | 199 | 200-217 | 0.65 | hydrolase, glucan 1,3-beta-glucosidase | 196-355 | 39.5 | 18 | YES | | | | | Group 3 |
| | | | | | | | lyase, tailspike gp27 | 339-664 | 24.9 | 22 | YES | | | | | |
| | | | | | | | hydrolase, exo-poly-alpha-d-galacturonosidase | 181-421 | 19 | 14 | YES | | | | | |
| | | | | | | | hydrolase, pectate lyase | 196-481 | 18.9 | 9 | YES | | | | | |
| vBKpMFBKp24_310 | QQV92016.1 | hypothetical protein | 679 | 193 | 194-212 | 0.62 | hydrolase, tail fiber | 109-579 | 99.8 | 23 | YES | | | | | Group 3 |
| | | | | | | | viral protein, hydrolase, tailspike protein | 110-673 | 100 | 18 | YES | | | | | |
| | | | | | | | hydrolase, alpha-1,3-glucanase | 57-415 | 98.9 | 19 | YES | | | | | |
| vBKpMFBKp24_313 | QQV92030.1 | hypothetical protein | 553 | 51 | 52-60 | 0.67 | hydrolase | 4-541 | 98.3 | 15 | YES | | | | | Group 4 |
| | | | | | | | hydrolase, xylosidase | 4-411 | 97.6 | 14 | YES | | | | | |
| | | | | | | | lyase, rhamnogalacturonan lyase | 4-343 | 97.9 | 12 | YES | | | | | |
| | | | | | | | single-stranded right-handed beta-helix, pectin lyase-like | 8-221 | 94 | 20 | YES | | | | | |
| | | | | | | | lyase, alginate lyase | 4-243 | 91.1 | 17 | YES | | | | | |
